# Long-term *in vivo* imaging of *Drosophila* larvae

**DOI:** 10.1101/744383

**Authors:** Parisa Kakanj, Sabine A. Eming, Linda Partridge, Maria Leptin

## Abstract

The *Drosophila* larva has been used to investigate many processes in cell biology, including morphogenesis, physiology, responses to drugs and new therapeutic compounds. Despite its enormous potential as a model system, it has technical limitations in cases where longer-term live imaging is necessary, because of the lack of efficient methods for immobilising larvae for extended periods. We describe here a simple procedure for anaesthetisation and long-term *in vivo* imaging of the epidermis and other larval organs including gut, imaginal discs, neurons, fat body, tracheae and haemocytes, and show a procedure for probing cell properties by laser ablation. We include a survey of different anaesthetics, showing that short exposure to diethyl ether is the most effective for long-term immobilisation of larvae. This method does not require specific expertise beyond basic *Drosophila* genetics and husbandry, and confocal microscopy. It enables high-resolution studies of many systemic and sub-cellular processes in larvae.

## Introduction

The fruit fly *Drosophila* is one of the most widely used multicellular organisms to study general biological processes. Originally the adult fly was used to work out the rules of chromosomal inheritance, then the embryo became the subject for the study of the genes that direct development. More recently, *Drosophila* has been used to study medically relevant problems, ranging from wound healing to cancer, neurodegenerative diseases, metabolism and ageing^1–16^. The available genetic techniques allow gene manipulation (both gain- and loss-of-function) with high spatial specificity and temporal control.

There have been very good reasons for adopting fly larvae and pupae as experimental systems. For example, many gene functions are not required, and many important developmental and physiological processes do not occur in the embryo and can therefore not be studied. Processes that can be better or exclusively studied at larval stages in development include growth control, organ innervation, memory formation and behaviour, tumourigenesis and others.

Investigation of many scientific questions is aided by live imaging of the processes under study. While all developmental stages from embryo to larva, pupa and adult fly, have been exploited for experimental analysis, the subject for live imaging has in the past mainly been the embryo. It was one of the early model systems used for live imaging, even before the introduction of GFP, largely because it is translucent and immobile, two important prerequisites for long-term live imaging^17, 18^.

The later stages in the *Drosophila* life cycle are also amenable to live imaging but pose greater technical and physiological challenges^19–22^. The pupa has been used for live imaging of a variety of cellular events^23–28^, but dissection is cumbersome, and is therefore the stage of choice only in especially justified cases.

The larva is suitable for imaging, but a major obstacle for long-term imaging has been the fact that larvae continuously move and are difficult to immobilize for extended periods.

Methods used in the past for imaging larvae relied on immobilisation by mechanical means or treatment with anaesthetics^21, 29–33^. Drawbacks of these highly useful methods include undesired side effects, such as stress through mechanical pressure, low viability of the larvae, and the restriction of immobilisation to short periods.

We have developed a method that overcomes these problems and allows immobilisation of larvae over extended periods^34^. We established laser ablation methods to induce defined wounds in the epidermis and to analyse tension in the larval epidermis while avoiding damage to other organs ^34, 35^. We have used the method to study aspects of signal transduction during epidermal wound healing^34^. As we show here, the method also allows imaging of internal organs like gut, trachea, imaginal discs, neurons, muscles and haemocytes and is suitable to study many aspects of cellular and subcellular events.

### Advantages of *Drosophila* larvae as a model system

The larva has all of the general advantages of *Drosophila* as a model system. The transgenic tools like Gal4 drivers, fluorescently tagged proteins and other markers, which were originally developed and established for studies in embryos can also be used for larvae. In addition, larvae are resilient to heat-shock or UV irradiation, the methods used to induce mosaic clones and to control temperature sensitive protein function and optogenetic tools^34, 36–38^.

In the embryo, many proteins and RNAs are provided maternally and persist throughout embryonic life, which can interfere with early genetic loss-of-function studies^39–43^. In the larva, most genes can be knocked down very effectively, since most maternally contributed proteins that may mask gene functions in the embryo have been depleted by this stage.

The larva is translucent and can be imaged without dissection. Many tissues are polyploid, as a result of which the cells are very large. This makes them ideal for imaging and for the study of subcellular processes; they are easily accessible for single-cell manipulations without risking damage to neighbouring cells or other organs^34^.

A range of interesting physiological processes occur during the larval stages of the life cycle. Most importantly, this is the main period of growth of the animal, during which the final body size of the adult fly is established. Growth is controlled by conserved signalling pathways, including the nutrient sensing pathways and practically all of the oncogenic pathways^44–49^. Growth can occur by cell proliferation, for example in imaginal discs and the brain, or through cell growth based on endoreduplication, as in the epidermis or salivary gland^50^. The patterning of differential growth is controlled by conserved signalling pathways and developmental genes (e.g. Dpp/BMP, Wnt, Hh and EGF pathway) and the larva and its organs present an excellent system to analyse stage- and tissue specific differential organ growth^51, 52^. For these reasons, the *Drosophila* larva has also become a well-established system for cancer research to study tumour initiation, progression and metastasis in different organs^53–56^.

Finally, it is easy to treat larvae with drugs, such as rapamycin, chloroquine, bafilomycin A1, wormatin, blebbistatin and many others, which can simply be administered by including them in the food^34, 57–59^. The larva therefore represents an attractive *in vivo* model both for fundamental and medically relevant questions.

### Development and validation of the protocol for long-term *in vivo* imaging of *Drosophila* larvae

Based on classic methods originally used to anaesthetise adult flies, M. Galko introduced the use of an anaesthetic chamber for brief immobilisations of larvae to induce epidermal wounds without subsequent live imaging^60, 61^.

We adapted this anaesthetic chamber and developed the protocol described here for immobilizing larvae for extended periods of live imaging^34^.

We have successfully used it to create epidermal wounds by laser, followed by long-term 3D *in vivo* confocal imaging (up to 8 hours) to document subcellular signalling and dynamics of wound healing^34^. We have also used the method for laser cuts and quantitative analysis of cell junctional tension in the epidermis^35^.

### Comparison with other methods

*Drosophila* larvae have in the past been immobilised for live imaging either by mechanical methods or by anaesthetics.

One method of mechanical immobilisation uses pressure on the larva through water capillary force^29^. A single larva is pressed between two coverslips with the help of a drop of water, which allows immobilisation for short-term live imaging (5-10min). Keeping larvae under pressure for more than 15 min kills them^29^.

In another method, larvae are immobilized in a microfluidic chip^30, 62^. Two microfluidic devices were designed, one for short-term imaging (up to 1 hour), the other for long-term imaging (up to 12 hours) of early 3rd instar larvae. In both devices, a constant vacuum maintains a strong seal between the chip, larva and coverslip interface to immobilise the larva. For long-term immobilisation a pulsed CO_2_/air mixture is supplied under moderate pressure to the microfluidic chip. Because these methods are based on mechanical pressure and squeezing of the larvae, they may induce acute and chronic stress responses and may affect signalling pathways, gene expression and changes in the mechanical tension and shape of the tissues. Supplying CO_2_ could also interfere with aspects of systemic physiology.

The main anaesthetic of choice for larvae has been desflurane^21, 31–33^. Anaesthetising with desflurane is more efficient than mechanical methods, and larvae survive better^21, 31^. However, one difficulty is that desflurane evaporates rapidly^21, 31^. As soon as it evaporates, the larvae start to move, necessitating repeated treatments for longer observations. The most complex and critical step in this method is applying desflurane during imaging.

Fuger et al. developed a chamber for continuous desflurane exposure during imaging^21^. The chamber is composed of two complex parts, a vaporizer/anaesthetisation device and an imaging chamber^21, 63, 64^. In this approach, a single larva is repeatedly exposed to desflurane for 5-10 min periods with recovery intervals of 5 min, with the cycle repeated up to 10 times. Heemskerk et al. developed a chamber for desflurane exposure with a simpler design for multiple rounds of anaesthetisation distributed over several hours (4 times over 6 hours)^31^.

Neither method allows continuous imaging over long periods because the larvae only survived short doses of the anaesthetic. Each round of anaesthetisation requires around 15-30 min preparation, which includes washing the larvae, a short pre-anaesthetisation to orient the larvae in the chamber, exposure to desflurane, placing the chamber under the microscope, focusing and adjusting of software. Both desflurane chambers can be used only on inverted microscopes because of their height^21, 31^.

All of these methods were designed for single larvae, which are individually immobilized and imaged. Thus, they are time consuming and demand extended usage of microscopes, which means they are not ideal for collecting data for statistical analysis and forward or reverse screening.

In a recent study Duygu et al. developed a simpler protocol for immobilizing up to 5 larvae at a time, without the use of manipulation devices, vaporizers or imaging chambers^33^. A comparison of desflurane and chloroform showed that desflurane arrested the heart beat in the first minutes, but this was followed by rapid recovery. Chloroform provided more rapid anaesthesia but slower recovery. The method is appropriate for brief, intermittent periods of *in vivo* imaging.

The protocol we describe here uses diethyl ether for anaesthetisation. A single and short-term exposure of the larvae to diethyl ether (3-4.5 min, depending on larval stage, see table 1 and 2) is sufficient to keep the larvae immobile for up to 8 hours. An additional advantage of this method is that up to 30 larvae can be immobilised and mounted for microscopy simultaneously and imaged in parallel.

**Fig. 1.**
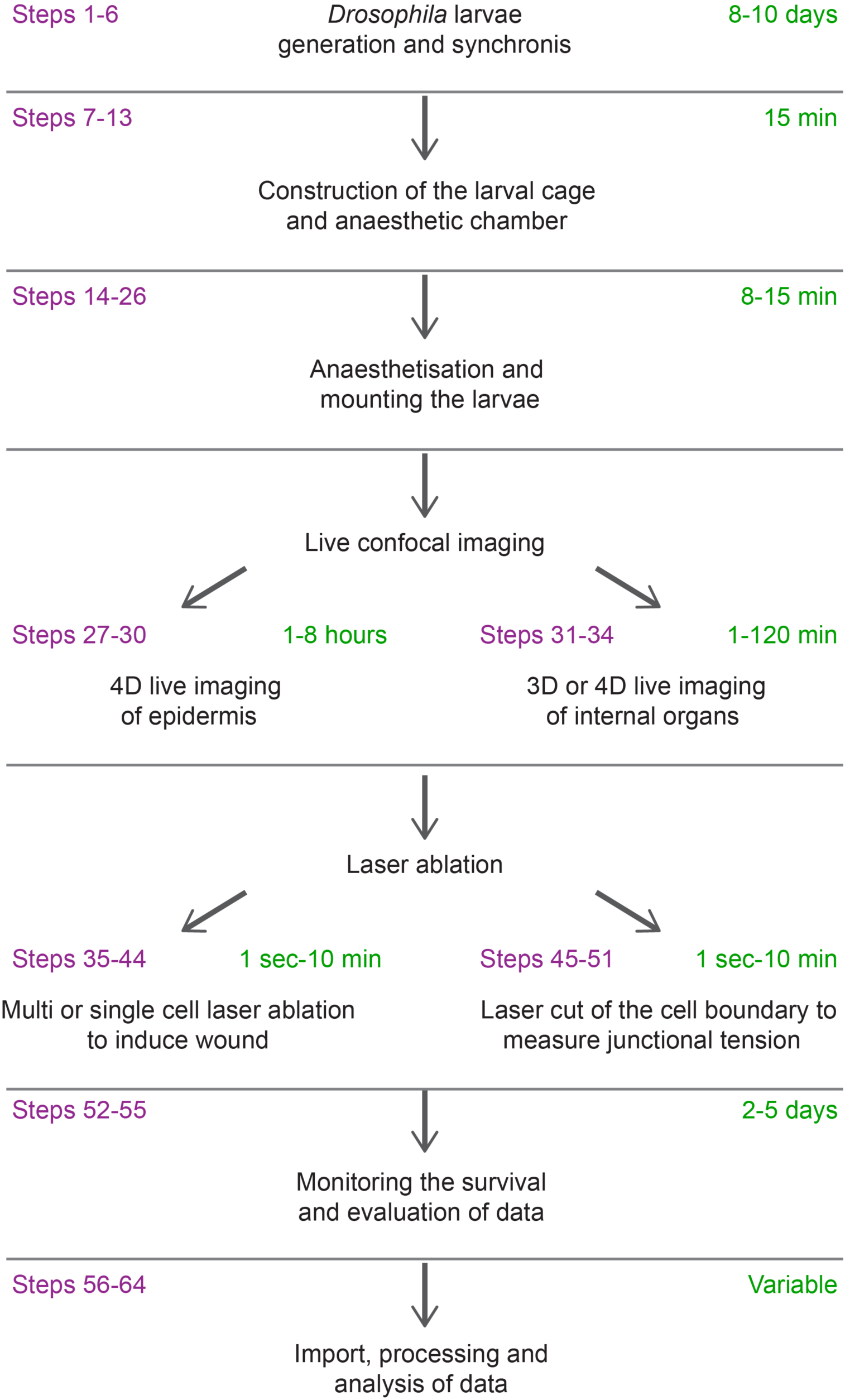
Outline of the experimental procedure. Experimental steps (magenta) and the anticipated duration (green) required for each step in minutes (min) are summarized in this image.

**Fig. 2.**
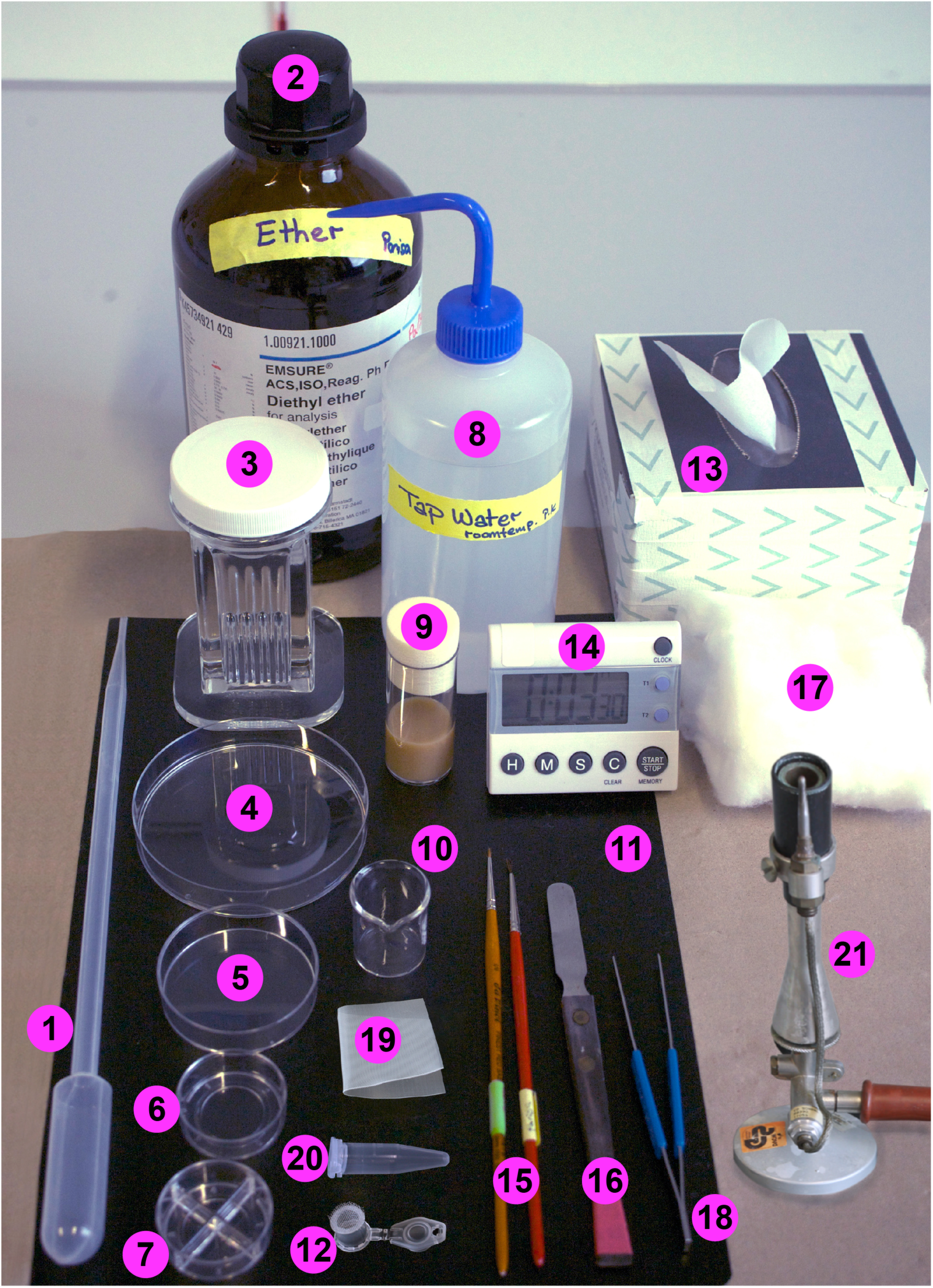
Overview of equipment required to anaesthetise larvae. (**1**) large plastic pipette, (**2**) diethyl ether, (**3**) glass Wheaton Coplin staining jar with PP screw cap, (4) culture dish (big), (**5**) culture dish (small), (**6**) culture dish for imaging with one compartment, (**7**) culture dish for imaging with four compartments, (**8**) tap water, (**9**) empty food vial to raise the larvae after imaging, (**10**) 10 ml glass beaker, (**11**) black nylon sheet, (**12**) self-made larval cage with lid (see **Fig. 3**), (**13**) paper napkin, (**14**) timer, (**15**) two paint brushes with real hair, size 1 and size 2/0, (**16**) spatula, (**17**) cotton wool, (**18**) filter forceps, (**19**) nylon mesh, 300-400 µm pore size, (**20**) 1.5 ml microcentrifuge (Eppendorf Safe-Lock tubes), (**21**) Bunsen burner. The numbers on this image correspond to the number sin the equipment section (for details see the **Equipment** section).

**Table 1.**
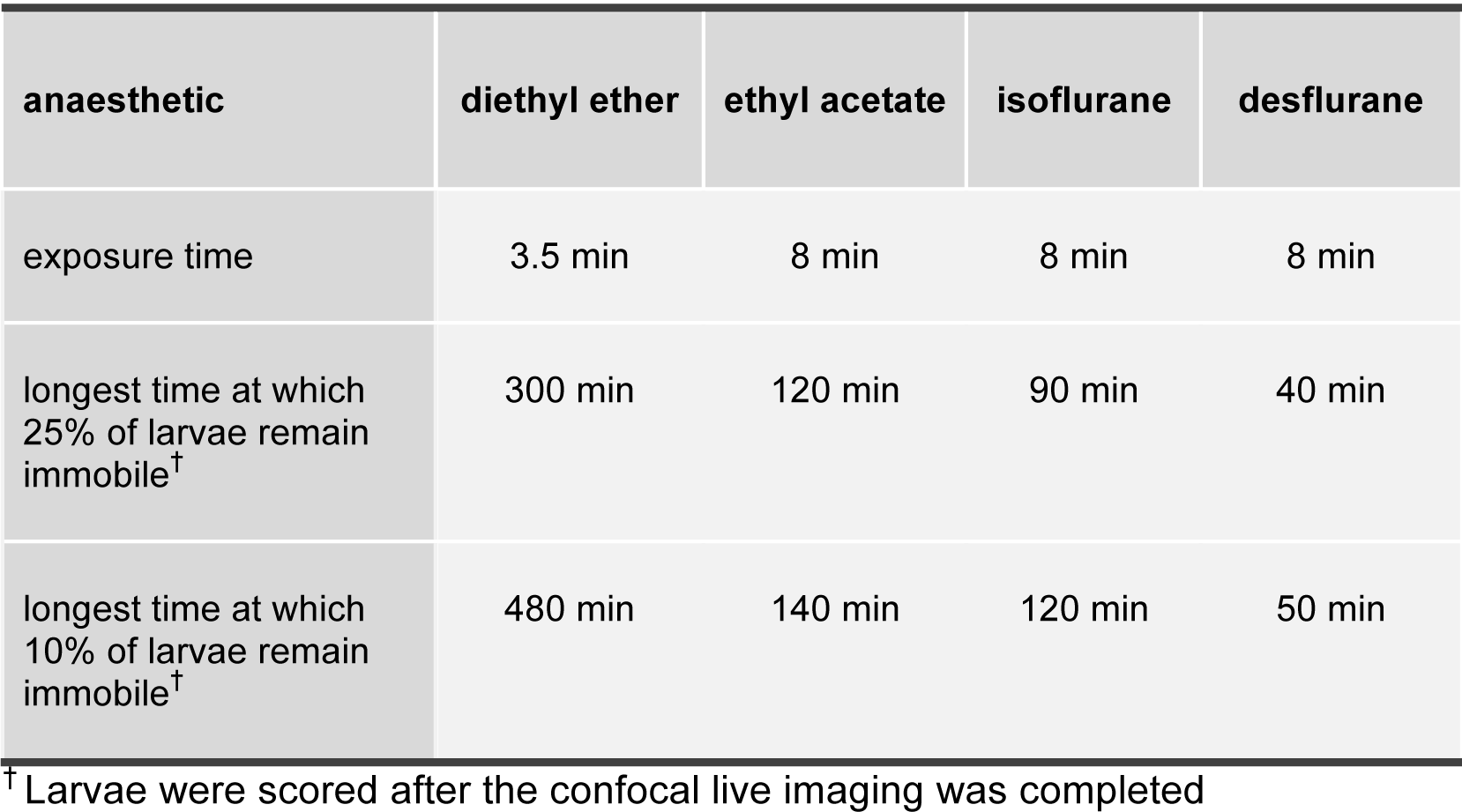
Comparison of different anaesthetics for immobilisation of larvae.

| anaesthetic | diethyl ether | ethyl acetate | isoflurane | desflurane |
| --- | --- | --- | --- | --- |
| exposure time | 3.5 min | 8 min | 8 min | 8 min |
| longest time at which 25% of larvae remain immobile<sup>†</sup> | 300 min | 120 min | 90 min | 40 min |
| longest time at which 10% of larvae remain immobile<sup>†</sup> | 480 min | 140 min | 120 min | 50 min |

<sup>†</sup> Larvae were scored after the confocal live imaging was completed

### Applications of the method

Long-term live imaging of *Drosophila* larvae can be used to visualize many organ systems: epidermis, neurons, trachea, muscle, fat body, gut, haemocytes, imaginal discs, and others. Any subcellular processes for which there are live imaging markers can be studied, e.g. calcium waves^34^, phagocytosis^34^, endocytosis, ER or Golgi vesicle trafficking, mitochondrial fusion and division, autophagy or cell death.

The same method can be used to obtain single images as a substitute for immunofluorescence on fixed specimens, as long as a fluorescent marker for the protein of interest is available. Anaesthetising and imaging 20-30 larvae in parallel takes 30-120 min, whereas immunofluorescent staining of larval organs 2-5 days and it is difficult to handle more than 20 larvae for each round of staining.

In addition to its high efficiency this protocol is fast, cheap and can easily be adapted to other chemical anaesthetics such as desflurane, isoflurane, ethyl acetate, as will be discussed below.

This method requires no special expertise or instrumentation. It is assumed that anyone wishing to apply it will already be proficient in *Drosophila* genetics and husbandry as well as general confocal microscopy.

### Limitations of long-term imaging

There are some limitations inherent to long-term imaging of larvae, irrespective of the method of immobilisation. During larval stages the animals normally feed constantly and grow rapidly. Long-term imaging induces two types of stress. First, feeding ceases and the larvae are starved. Indeed after 4-6 hours starvation, autophagy can be seen in most issues^65–67^. As observed previously in the fat body and muscles, autophagosomes first appear in the epidermis after 4.5 hours of imaging^34^. Thus, at least in the first 4.5 hours, this anaesthetisation method does not induce autophagy and can be used without concern.

The constant illumination during imaging can also cause stress. The laser beam concentrates energy locally to visualise fluorescent proteins, which can lead to phototoxicity and local heat stress. However, these problems apply to any long-term live imaging experiment in any cell type or organism.

### Overview of procedure and experimental design

We describe here a protocol for long-term *in vivo* confocal imaging of *Drosophila* larvae. Originally developed to study epidermal wound healing^34^, this now improved protocol can be applied to study any cellular or sub-cellular events in any organ, as outlined in detail in the **Procedure** section.

We also describe how the same equipment and chamber can be used to anaesthetise the larvae with other anaesthetics than diethyl ether and compare the effect of desflurane, isoflurane, ethyl acetate and diethyl ether on immobilisation and survival of the larvae (Table 1 and 2).

We provide a list of Gal4 drivers (Table 3), markers for visualising subcellular compartments and organelles (Table 4) and useful GFP protein traps, which we tested and found most useful for live imaging in larvae (Table 4).

The steps of the procedure are the following (Fig. 1):

- Selection and generation of optimal lines for live imaging (Steps 1-4)
- Staging and synchronizing larval growth (Steps 5-6)
- Construction of larval cage and anaesthetic chamber (Steps 7-13)
- Anaesthetisation of larvae (Steps 14–26)
- Microscope parameters for optimal long-term *in vivo* imaging of epidermis and internal organs in the larva (Steps 27–34)
- Multi- and single-cell laser ablation for wound healing studies (Step 35-44)
- Laser cutting of cell boundaries to measure junctional tension (Step 45-51)
- Monitoring the survival of the animals and evaluation of data for quantitative measurements (Steps 52-55)
- Import and processing of images (Steps 56-60)
- Analysis and quantification of wound closure and junctional tension (Step 61-64)

### Selection and generation of lines for live imaging (Steps 1-4)

The experimental expression of genes with the help of the Gal4/UAS system is a common and efficient method to mark tissues or cells of interest and to manipulate gene function in *Drosophila*^68^. The lab of M. Galko established *Drosophila* larvae for epidermal wound healing experiments and developed a number of useful tools for the larval epidermis^61, 69^. For imaging of epidermal wound healing, we took advantage of the existing tools and used two lines carrying Src-GFP to visualise the plasma membrane and DsRed2-Nuc to mark the nuclei in the epidermis of the larvae. The stocks also carry a Gal4 driver, either *A58-Gal4* or *e22c-Gal4,* that direct expression in the epidermis^61, 70^.

We recommend testing Gal4 and UAS lines for their specificity and efficiency as part of the design of the experiment, especially if they will be used for gain- and loss-of-function studies. We tested a range of Gal4 driver lines for their tissue specificity. None directed expression in only one larval tissue. The selected Gal4 drivers, fluorescent reporter constructs and GFP-traps we used are listed in tables 3 and 4. In the experiments to measure the recoil velocity of the cell cortex after laser cutting we visualised cell outlines with an endogenously GFP-tagged DE-cadherin^35, 71^.

For precise visualisation and analysis the expression of fluorescent markers should ideally be restricted to the tissue of interest so that the surrounding tissues are free of fluorescent proteins. Fluorescent signal from underlying organs, in particular the fat body, makes it difficult to detect the signal in the tissue of interest.

All red fluorescent markers (mCherry, RFP, Red2, Tomato) we have tested in epidermis and other organs, regardless of which proteins they tagged, showed cytosolic aggregates that were seen as very bright spots. These do not necessarily interfere with detecting the fusion protein proper (for example, when visualising actomyosin cables or FOXO shuttling)^34^ but may influence aspects of quantification, especially if small organelles or cytoplasmic vesicles are to be counted. Therefore, quantitative and qualitative analysis with mCherry, RFP, R d2, Tomato, etc. should be performed very carefully and critically.

To express constructs in the epidermis, four Gal4 drivers can be used: *e22c-Gal4*, *A58-Gal4, 69B-Gal4* and *da-Gal4*^61, 68, 70, 72^. The best marker constructs for cell outlines and membranes are UAS-Src-GFP (a myristoylated GFP fusion protein) and UAS-mCD8-GFP (a transmembrane GFP fusion protein). They can be imaged for long periods without much bleaching.

### Staging and synchronizing the larval growth (Steps 5-6)

If several larvae are to be imaged simultaneously or mutants are to be compared with control larvae, larval growth has to be synchronised to obtain larvae that are at the same developmental stage. This is not only helpful for the experimental design, but also necessary because different stages require different durations of exposure to the anaesthetic (Table 2).

**Table 2.** Time of exposure to anaesthetic for live imaging of larvae in different instar larval stages.

| desired imaging time | anaesthetic | L1 | early L2 | late L2 | early L3 | middle L3 |
| --- | --- | --- | --- | --- | --- | --- |
| 5 h | diethyl ether | - | - | 3 min | 3.5 min | 4 min |
| 7-8 h | diethyl ether | - | - | 3 min | 3.5 min | 4.5 min |
| 10-20 min | desflurane | 1 min | 2 min | 3 min | 3.5 min | 3.5 min |
| 30-50 min | desflurane | - | - | 6 min | 8 min | 8 min |
| 80-120 min | ethyl acetate | - | - | 6 min | 8 min | 8 min |
| 70-90 min | isoflurane | - | - | 6 min | 8 min | 8 min |

L: larval instar

### Construction of larval cage and anaesthetic chamber (Steps 7-13)

An overview of all materials required for this experiment is shown in **Fig. 2** and the experimental setup required to anaesthetise the larvae and mount them for confocal live imaging is demonstrated in **Figs. 3 to 5**. We used the design from the Galko lab^60^ for the larval cage (Fig. 3) and anaesthetic chamber (Fig. 4) that are easy, cheap and fast to construct.

**Fig. 3.**
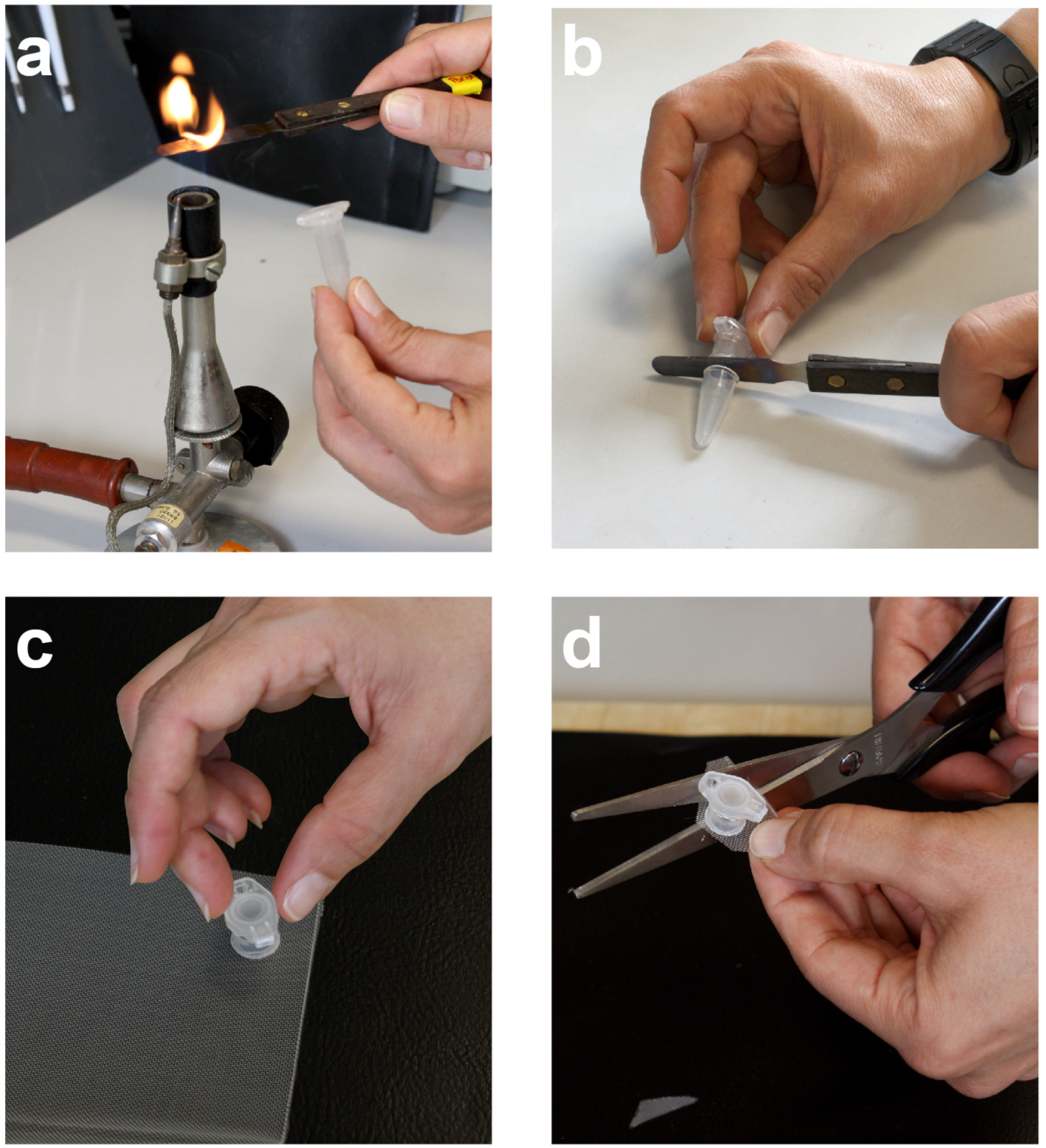
Construction of the larval cage. Image shows how to make a larval cage needed for anaesthetisation of larvae. For details see **step 9**.

**Fig. 4.**
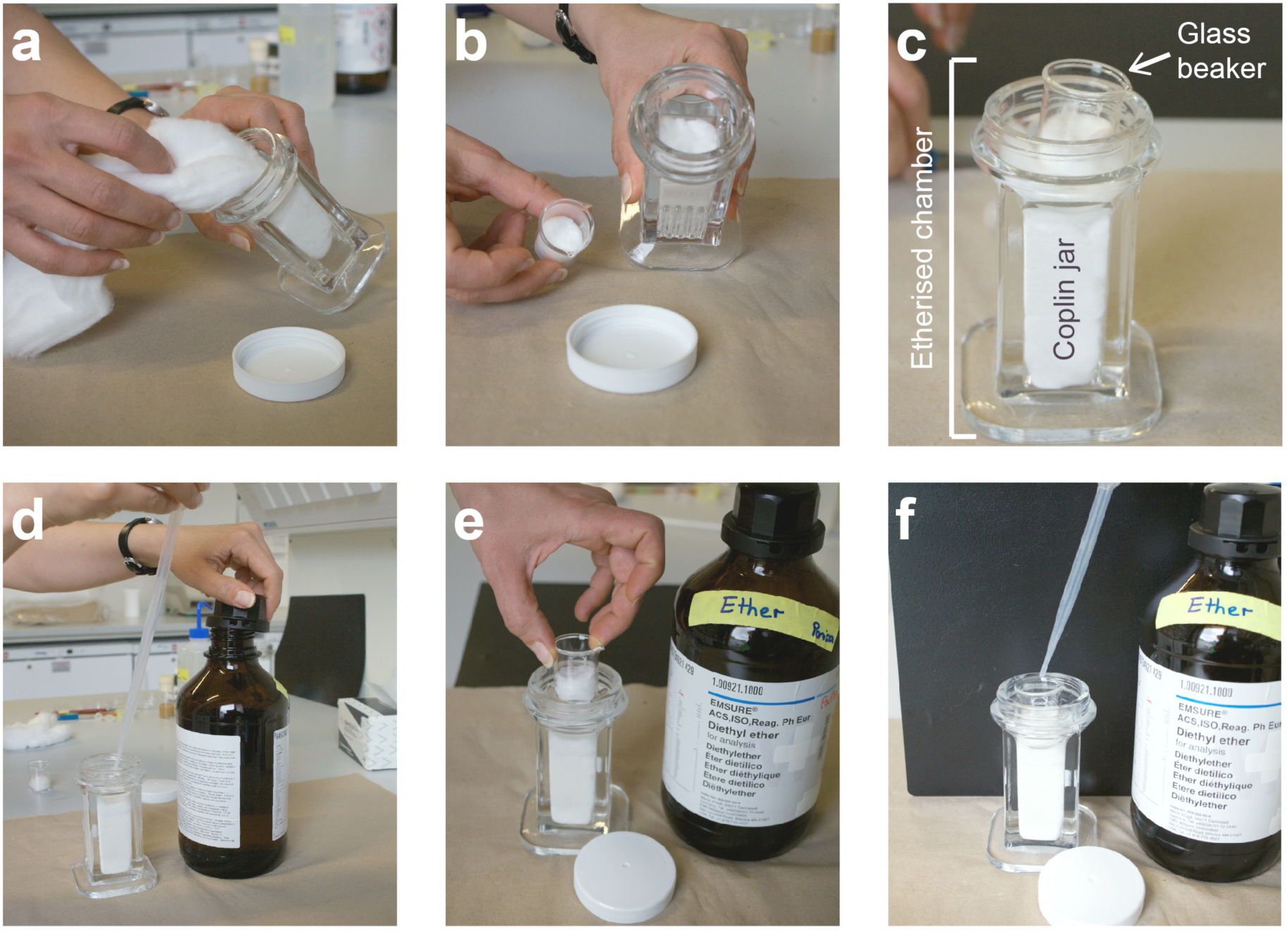
Construction of anaesthetisation chamber. This image shows the steps to construct an anaesthetisation chamber required for immobilisation of the larvae. For details see **step 10-16**.

**Fig. 5.**
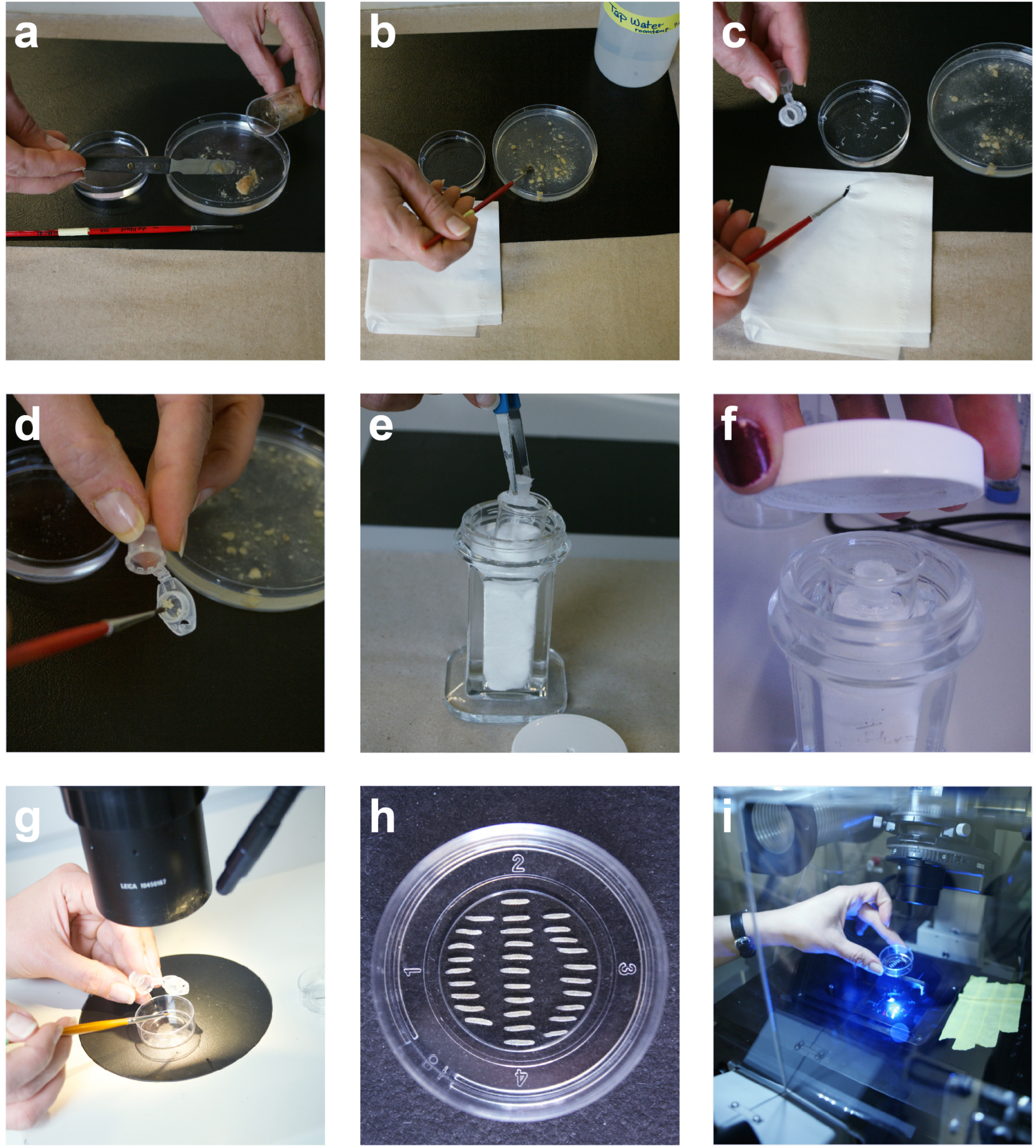
Anaesthetisation and mounting the larvae for live imaging. Sequence of steps to anaesthetise and mount the larvae for live imaging. (**a-c**) Sort, wash and dry the larvae. (**d**) Place larvae into the larval cage. (**e**) Place the larval cage inside the glass beaker. (**f**) Close the screw cap of the anaesthetic chamber and expose the larvae for 3-3.5 min (see table 2) to diethyl ether. (**g-h**) Transfer the anaesthetised larvae into an imaging dish, sort and orient them under the stereo microscope. (**i**) Now the larvae are ready for imaging. Place the imaging dish under the microscope. For details of the procedure see **steps 16-28**.

### Anaesthetisation of larvae (Steps 14–26)

The anaesthetics we have tested (diethyl ether, desflurane, isoflurane and ethyl acetate) are liquids and evaporate when placed into the chamber. Since we found diethyl ether to provide the longest anaesthetising effect it was used in most of our experiments. The larvae remained immobile for 7-8 h if they were exposed for 3-4,5 min to diethyl ether in the anaesthetising chamber (Table 1 and 2).

To examine the effects on viability of the experimental procedure itself, and of the immobilisation with four different anaesthetics we scored the further development and survival of treated individuals (Fig. 6). As a baseline control, early third instar larvae were washed and starved for 5 h in an imaging dish without water or nutrients. Experimental animals were either washed and immobilised with the different anaesthetics (diethyl ether, ethyl acetate, isoflurane or desflurane) for 3.5 min or subjected to this treatment and starved as well. Each experiment was done in triplicate, with 30 heterozygous *A58>Src-GFP, DsRed2-Nuc* (*w^1118^; +; A58-Gal4, Src-GFP, DsRed2-Nuc/+*) larvae in each sample. After washing and sorting only, 84% of the larvae survived to adult stages. If, in addition, they were starved for 5 hours, 80% of the larvae survived until puparation and 71% until eclosion of the adult. This survival rate was not affected further by anaesthetisation with diethyl ether. After washing and exposure to diethyl ether for 3.5 min without subsequent starvation, 78% of the larvae survived until puparation and 67% eclosed as adult flies. With the combination of all manipulations - washing, sorting, diethyl ether exposure for 3.5 min and 5 hours starvation - 64% of the larvae survived to the pupal stage and 60% to adult flies.

**Fig. 6.**
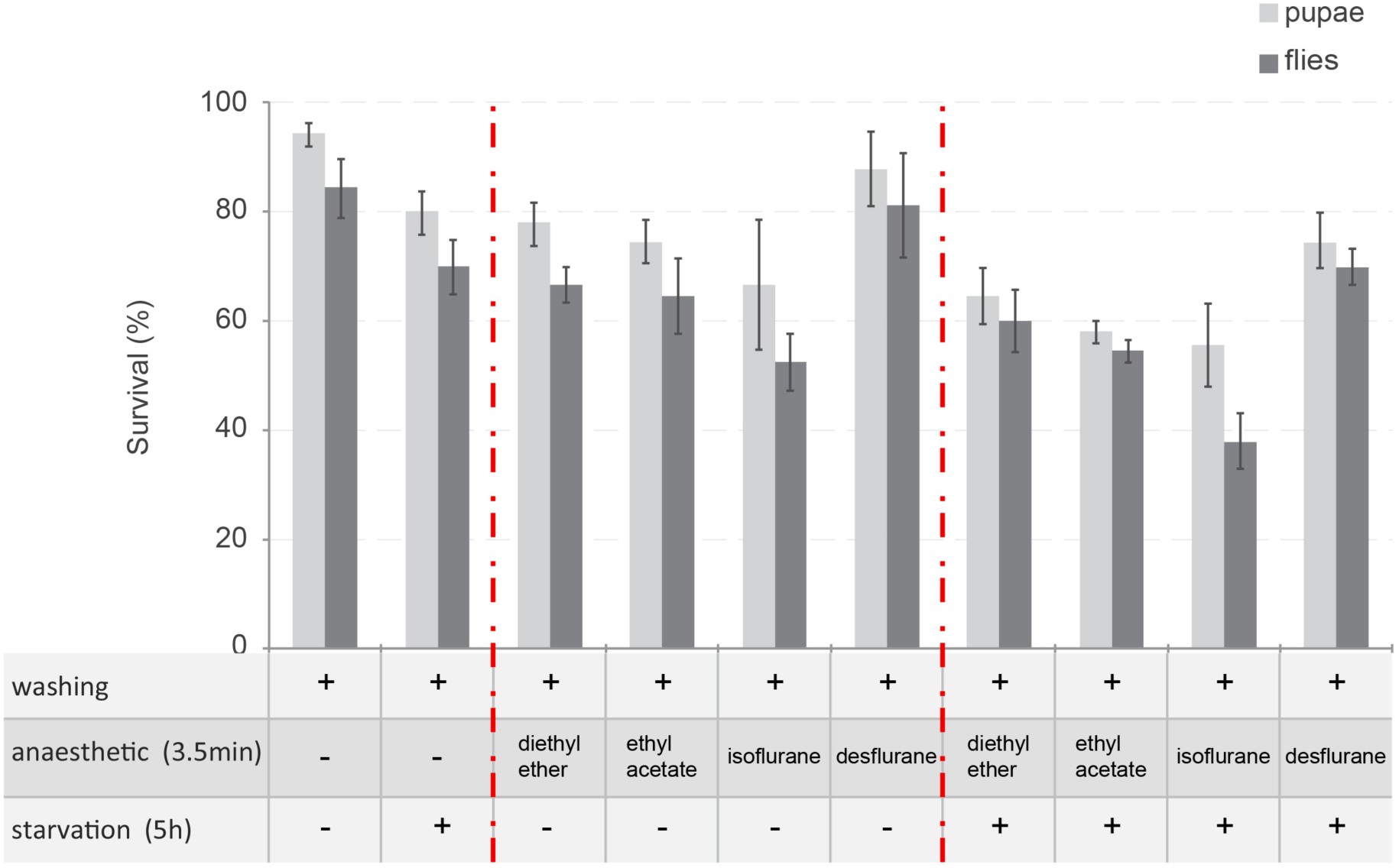
Effect of different anaesthetics on larval survival. The graph shows the proportions of L3 larvae that pupate (light grey) and develop to adult flies (dark grey) after of washing, 3.5 min exposure to anaesthetic (diethyl ether, ethyl acetate, isoflurane or desflurane) and 5 hours starvation, which corresponds to 5 hours of imaging without nutrient. Average number of survivors as pupae and adult flies for each condition tested. The experiments were done in triplicate, each with 30 heterozygous larvae for each condition. Data are given as mean ± s.e.m. Transgene genotypes: *A58>Src-GFP, DsRed2-Nuc (w^1118^; +; A58-Gal4, UAS-Src-GFP, UAS-DsRed2-Nuc/+*).

We also tested the effects of other anaesthetics on viability, both with and without long-term starvation (Fig. 6). Desflurane gives slightly better survival rates than diethyl ether, with or without starvation. Ethyl acetate is equal and isoflurane is worse than diethyl ether. Of all the anaesthetics, we found isoflurane to be the most toxic for *Drosophila* larvae.

Unlike diethyl ether, 3.5 min exposure to desflurane, isoflurane or ethyl acetate is not sufficient to anaesthetise larvae for long-term live imaging (Table 1 and 2). To immobilise them for more than 10-20 min, larvae have to be exposed for 8 min to desflurane, isoflurane or ethyl acetate. We have recorded the maximal imaging times during which at least 25-10% of the larvae remained immobile (Table 1). With the same exposure time (3.5 min) diethyl ether provided the longest (7-8 h) and desflurane the shortest (10-20 min) narcotising effect (Table 2). Thus, for long-term imaging of the larvae, diethyl ether is the most suitable anaesthetic, and more efficient than any other published treatment.

### Microscope set-up for optimal long-term *in vivo* imaging of epidermis and internal organs (Steps 27–34)

Before anaesthetising larvae and starting an experiment, it is important that the microscope is completely set up, with all parameters entered and ready for imagining.

For the long- and short-term live image acquisition of epidermis and other organs, a spinning disk confocal microscope is best suited for fast imaging and minimising photobleaching. For any given application, the total exposure time, laser power, frequency of time intervals, objective, number of sections per *z* stack, and number of fluorescence channels acquired have to be optimized empirically to achieve the desired balance between speed of imaging, image quality and photo-bleaching. The parameters used for long-term live imaging of larval epidermis in our own experiments are described in the **Procedure** section below. To trace fast subcellular processes within the epidermis (e.g. calcium flashes or vesicle trafficking), we used faster laser scanning (see **Procedure** section).

Live imaging of internal organs in larvae is not trivial. Although the anaesthetic stops somatic muscle movement, the peristaltic gut movement and heartbeat continue, and their movement is indirectly transmitted to other internal organs. However, with high-speed imaging at a rate that is considerably faster than the speed of internal organ movements, it is possible to image imaginal discs, tracheae, gut, muscle, neurons, fat body and haemocytes (Fig. 7, Supplementary Video 1-11). The microscope setting for imaging internal organs is described in the **Procedure** section below.

**Fig. 7.**
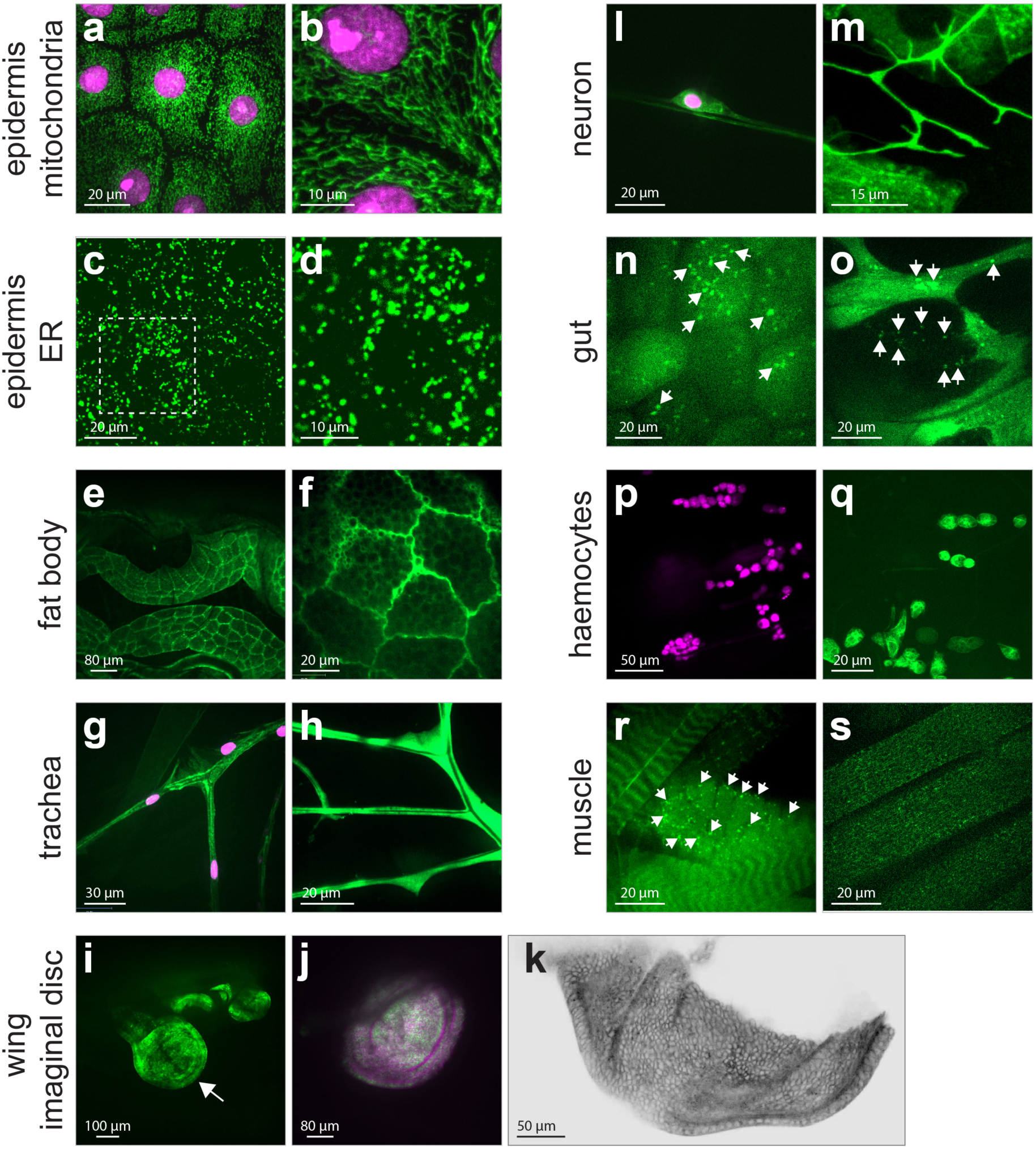
Live imaging of inner organs and subcellular organelles. (**a**-**b**) mitochondria (green) and nuclei (magenta) in the dorsal epidermis of larvae expressing *A58>mito-GFP; DsRed2-Nuc*, (**b**) higher magnification of epidermal mitochondria during wound healing, corresponds to **supplementary Video 1,** (**c**-**d**) ER (green) in the dorso-lateral epidermis of larvae, *e22c>; RFP-KDEL*, (**d**) higher magnification of area marked in (**c**), corresponds to **supplementary Video 2**, (**e**-**f**) fat body (green), *c7>mCD8-GFP*, (**f**) higher magnification, corresponds to **supplementary Video 3**, (**g**) trachea (green) and nuclei (magenta), *e22c>Src-GFP,DsRed2-Nuc*, (**h**) trachea (green), *btl>GFP*, corresponds to **supplementary Video 4,** (**i**) entire wing imaginal disc (green, white arrow)*, esg>GFP*, (**j**) *wing pouch region* (green), nuclei (magenta), *Pdm2>mCD8-GFP; DsRed2-Nuc,* (**k**) posterior compartment of wing imaginal disc (grey), *hh>mCD8-GFP*, **i** and **k** correspond to **supplementary Video 5,** (**l**) peripheral neurons (green) with nucleus (magenta), *Nub>mCD8-GFP; DsRed2-Nuc,* **(m)** peripheral neurons (green), *elav>mCD8-GFP*, corresponds to **supplementary Video 6,** (**n**) Midgut enterocytes (green), (**o**) midgut interstitial cells (green), *NP1>GFP-Atg8a; foxo*, (**n-o**) high level of FOXO in enterocytes and interstitial cells induce autophagy (green dots represent autophagosomes, arrows), **n** and **o** correspond to **supplementary Video 8 and 9, (p)** haemocytes (magenta), *Hml>dsRed*, **(q)** haemocytes (green), *Jupiter-GFP* trap, **p** and **q** correspond to **supplementary Video 10,** (**r**) muscle (green), *MHC>GFP-Atg8a; foxo*, overexpression of FOXO in muscle induces autophagy (green dots represent autophagosoms, arrows), (**s**) muscle (green), *Nrg-GFP trap*, **r** and **s** correspond to **supplementary Video 11. (a-b, g-h, m, p-q**) Projections of the epidermis from a time-lapse series. Each frame is a merger of (**a-b**) 57 or (**g-h, m, p-q**) 21 planes spaced 0.28 µm apart. (**c-f, i-l, n-o, r-s**) Single plane images from time-lapse series of different inner organs and subcellular organelles of L3 larvae. Transgene genotypes of larvae: (**a**, **b**) *w^1118^; UAS-mito-HA-GFP/+; A58-Gal4, DsRed2-Nuc/+*, (**c**, **d**) *w^1118^; e22c-Gal4/+; UASp-RFP.KDEL/+*, (**e**, **f**) *w^1118^; c7-Gal4/+; UAS-mCD8-GFP/+*, (**g**) *w^1118^; e22c-Gal4, UAS-Src-GFP, UAS-DsRed2-Nuc/+; +*, (**h**) *w^1118^; btl-Gal4, UAS-GFP; +*, (**i**) *w^1118^; esg-Gal4, UAS-GFP/+; +*, (**j**) *w^1118^; UAS-mCD8-GFP/+; Pdm2-Gal4/UAS-DsRed2-Nuc*, (**k**) *w^1118^; +; hh-Gal4, UAS-mCD8-GFP/+*, (**l**) *w^1118^; Nub-Gal4/UAS-mCD8-GFP/+; UAS-DsRed2-Nuc/+*, (**m**) *w^1118^, elave-Gal4/UAS-mCD8-GFP; UAS-mCD8-GFP/+; UAS-mCD8-GFP/+*, (**n**, **o**) *w^1118^; NP1-Gal4/UAS-GFP-Atg8a; UAS-foxo/+*, (**p**) *w^1118^; Hml-Gal4/+; UAS-dsRed/+*, (**q**) *w^1118^; +; Jupiter-GFP*, (**r**) *w^1118^; MHC-Gal4/UAS-GFP-Atg8a; UAS-foxo/+*, and (**s**) *w^1118^, Ngr-GFP; +; +*.

### Multi- and single-cell laser ablation for wound healing studies (Step 35-44)

The larval epidermis is an attractive model to study many cellular processes. Like simple epithelia in mammals, the larval epidermis is a monolayered epithelium with a basal lamina. On the outside it is covered with an apically secreted cuticle (Fig. 8)^34, 73^. In the case of epidermal wound healing, the absence of epidermal tissue remodelling (rearrangement or morphogenesis) is an additional advantage.

**Fig. 8.**
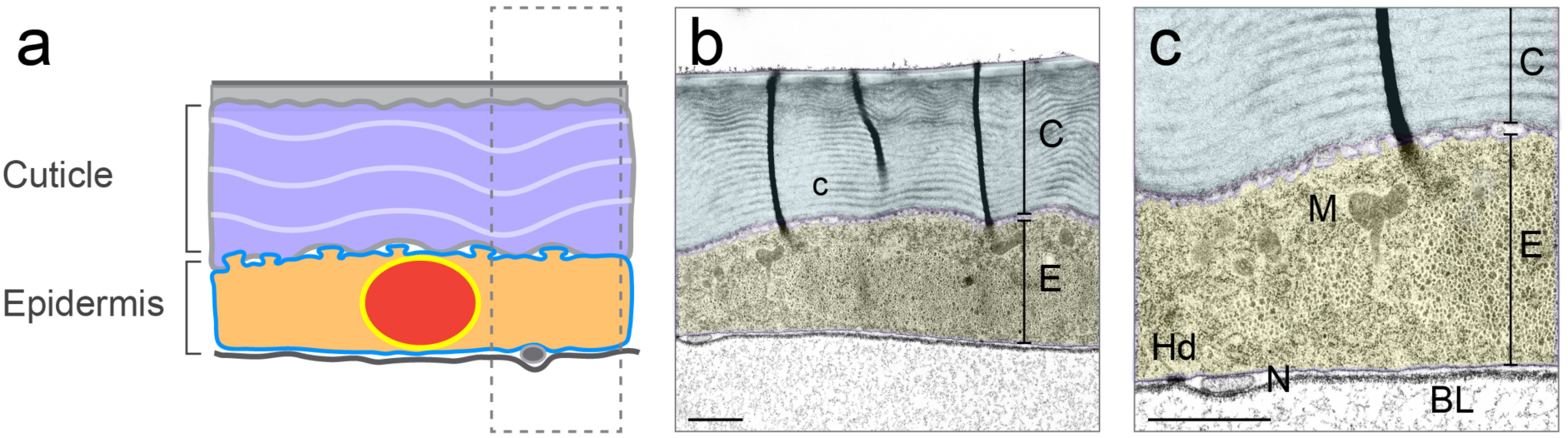
Epidermis of *Drosophila* larva. (**a**) Schematic diagram illustrating the structure of the larval epidermis. (**b-c**) Transmission electron microscopic picture of cross section through larval epidermis. (**c**) Magnification of **b.** (**a-c**) Epidermal cell is marked in yellow and secreted cuticle marked in blue. The larvae used for transmission electron microscopy had the following genotype: *w^1118^; e22c-Gal4, UAS-Src-GFP, UAS-DsRed2-Nuc/+; +*. Scale bars: (**b-c**) 1 µm. BL: basal lamina; C: cuticle; E: epidermis; Hd: hemidesmosome; M: mitochondria; N: neuron (axon).

For optimal documentation of subcellular events immediately after cell wounding we used the same microscope and lens for laser ablation and imaging. The set up and all parameters and detailed experimental procedure are described in the **Procedure** section.

### Laser cutting to measure junctional tension (Step 45-51)

To analyse the tension along cell-cell junctions we performed laser ablations with the settings described in the **Procedure** section and imaged with the same equipment used for wound healing, as described below (in the **Procedure** section).

### Monitoring the survival of the animals and evaluation of data to obtain quantitative measurements (Steps 52-55)

For reliable comparisons and quantifications of cellular dynamics, all larvae included in the study must have the same fitness. Not all larvae survive the laser wounding and live imaging. It is an advantage to be able to recognize weak larvae early and exclude them from the experimentation and further observation. Some weak larvae can be recognized and excluded before the beginning of the experiment; others are visibly damaged during the course the experiment. The most stringent criterion for quality is survival to adulthood after the experiment.

In some cases, low vitality can be judged by visual inspection. One sign of fitness is the larval heartbeat. Larvae in which the heart is not beating should be excluded from the experiment. Another indication that larvae are not fit is a reduction of intensity or an aggregation of fluorescent markers or appearance of regular, particulate structures (Fig. 9). Larvae in which the fluorescent markers appear even slightly abnormal should also be excluded (Fig. 9b). Indications that larvae are dying during the experiment include: loss of the fluorescent marker (Fig. 9a, h), break-down of the plasma membrane (Fig. 9a, f, g, h), continuing expansion of the wound and disintegration of tissue (Fig. 9g, h), formation of stress fibres (Fig. 9f), nuclear envelope breakdown (Fig. 9d, h, i) or fluorescent markers in inappropriate subcellular locations (Fig. 9b, c).

**Fig. 9.**
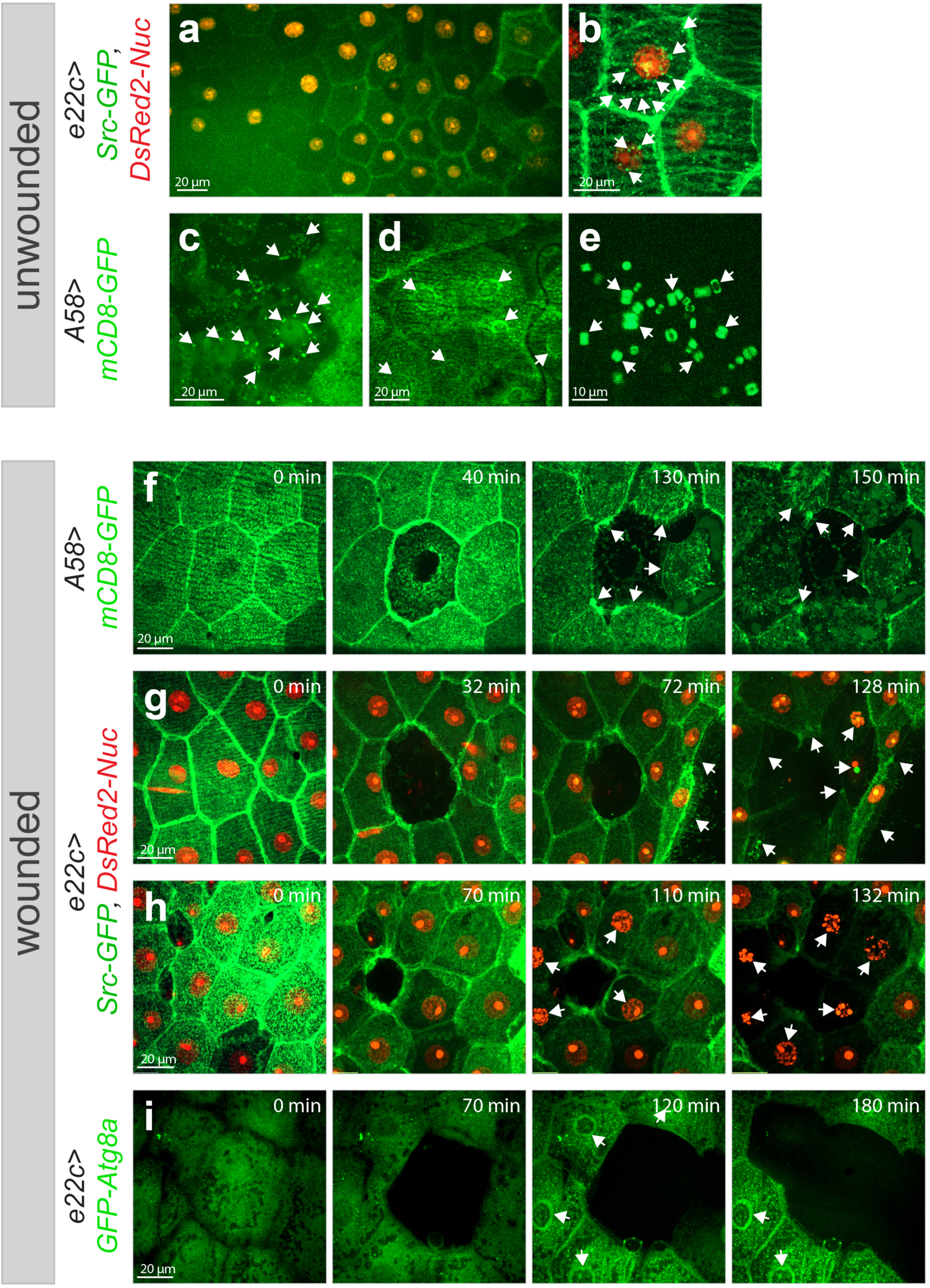
Signs of reduced larval vitality. Indications for larvae being unfit or dying (**a-e**) during imaging (without laser ablation) or (**f-i**) after laser ablation: (**a,h**) reduced intensity of fluorescent markers, (**b,c**) aggregation of fluorescent protein (arrows), (**e**) appearance of regular, particulate structures (arrows), (**f,g**) disintegration of tissue, break-down of the plasma membrane and continuous expansion of the wound, and formation of stress fibres (arrows) or (**d,i**) breakdown of nuclear envelope and appearance of cytoplasmic fluorescent protein in the nucleus or accumulation of fluorescent marker iat the nuclear envelop (arrows). (**a-e**) Projections of an image or (**f-i**) a time-lapse series of the wounded epidermis in third instar larvae. Each frame is a merge of 57 planes spaced 0.28 µm apart. Scale bars: (**a-d and f-i**) 20 µm and (**e**) 10 µm. Transgene genotypes: (**a-b, g-h**) *e22c>Src-GFP,DsRed2-Nuc (w^1118^; e22c-Gal4, UAS-Src-GFP, UAS-DsRed2-Nuc/+; +*), (**c-f**) *A58>mCD8-GFP (w^1118^; +; A58-Gal4, UAS-mCD8-GFP/+*) and (**i**) *e22c>GFP-mCherry-Atga8* (*w^1118^; e22c-Gal4/UASp-GFP-mCherry-Atg8a/+; +*) in image **i** only the green channel is shown.

### Importing and processing of images (Steps 56-60)

For visualisation and analysis, the imaging data have to be processed, converted and imported using appropriate software. We have mostly used the commercially available software Volocity (6.3.57 version). Other commonly used packages are ImageJ/Fiji (National Institutes of Health), an open source image-processing package, and Imaris (Bitplane) a commercially available software widely used for 4D data analysis.

### Analysis and quantification of wound closure and junctional tension (Step 61-64)

Depending on available software, the data can be analysed and quantified as raw data (e.g in Volocity) or as converted “ome.tiff” files (e.g. in Fiji). The details for quantification are described in the **Procedure** section.

## Materials

### Biological materials

- The Gal4 drivers, UAS-fluorescence reporter constructs and GFP-traps we used for imaging are listed in Tables 3 and 4. Further details are described in the **Procedure** section.

**Table 3.**
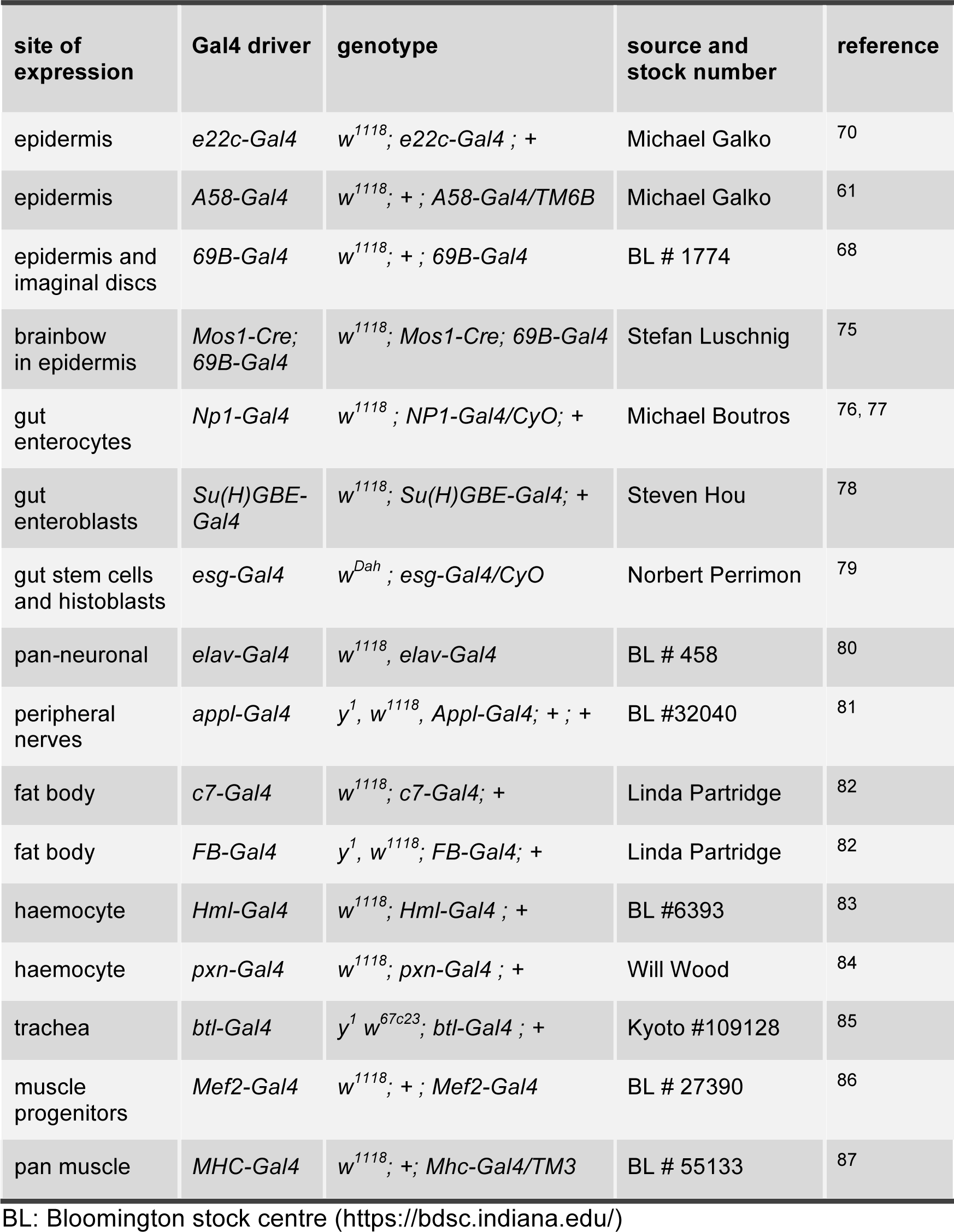
Gal4 lines for imaging various organs in *Drosophila* Larvae.

**Table 4.**
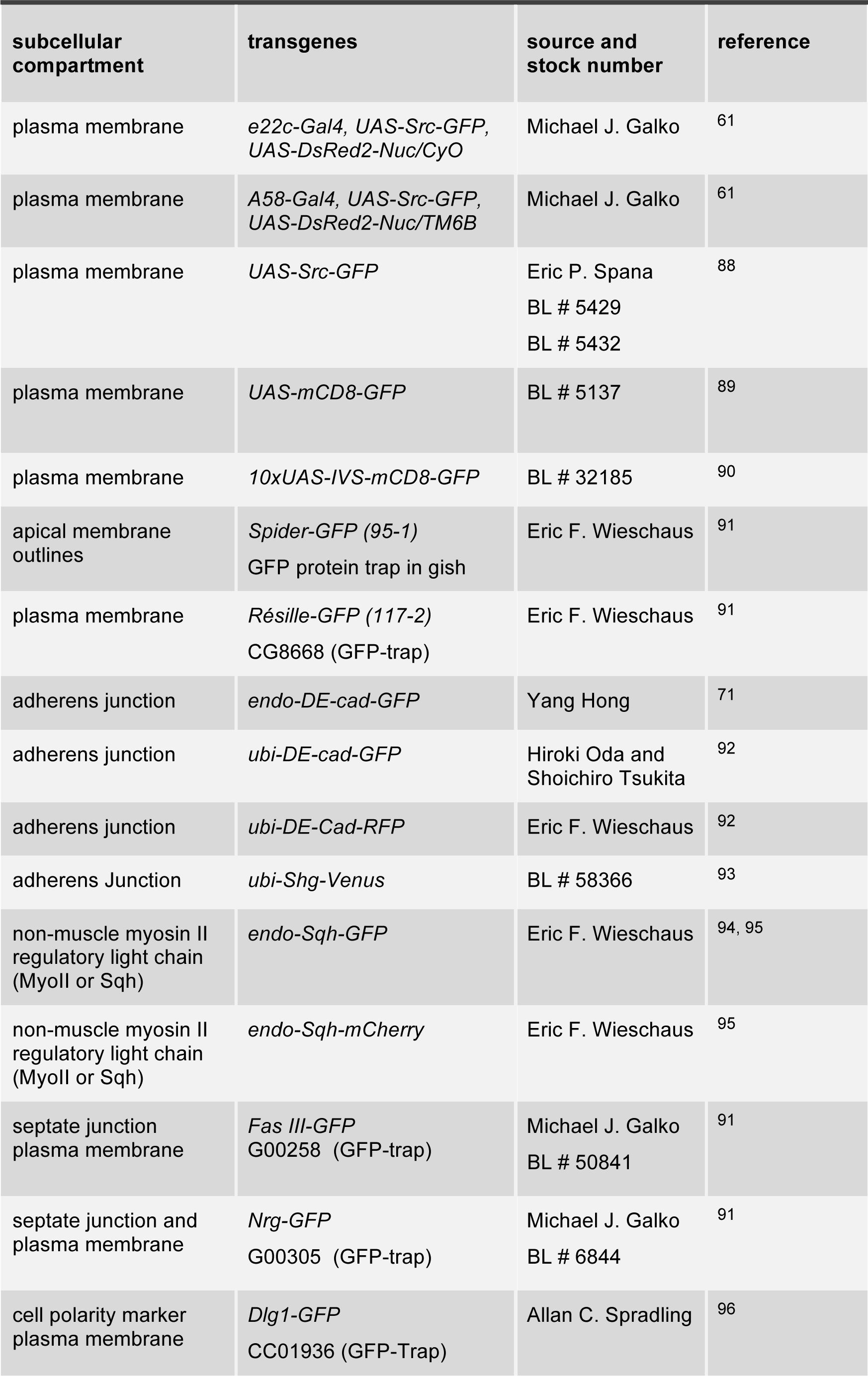
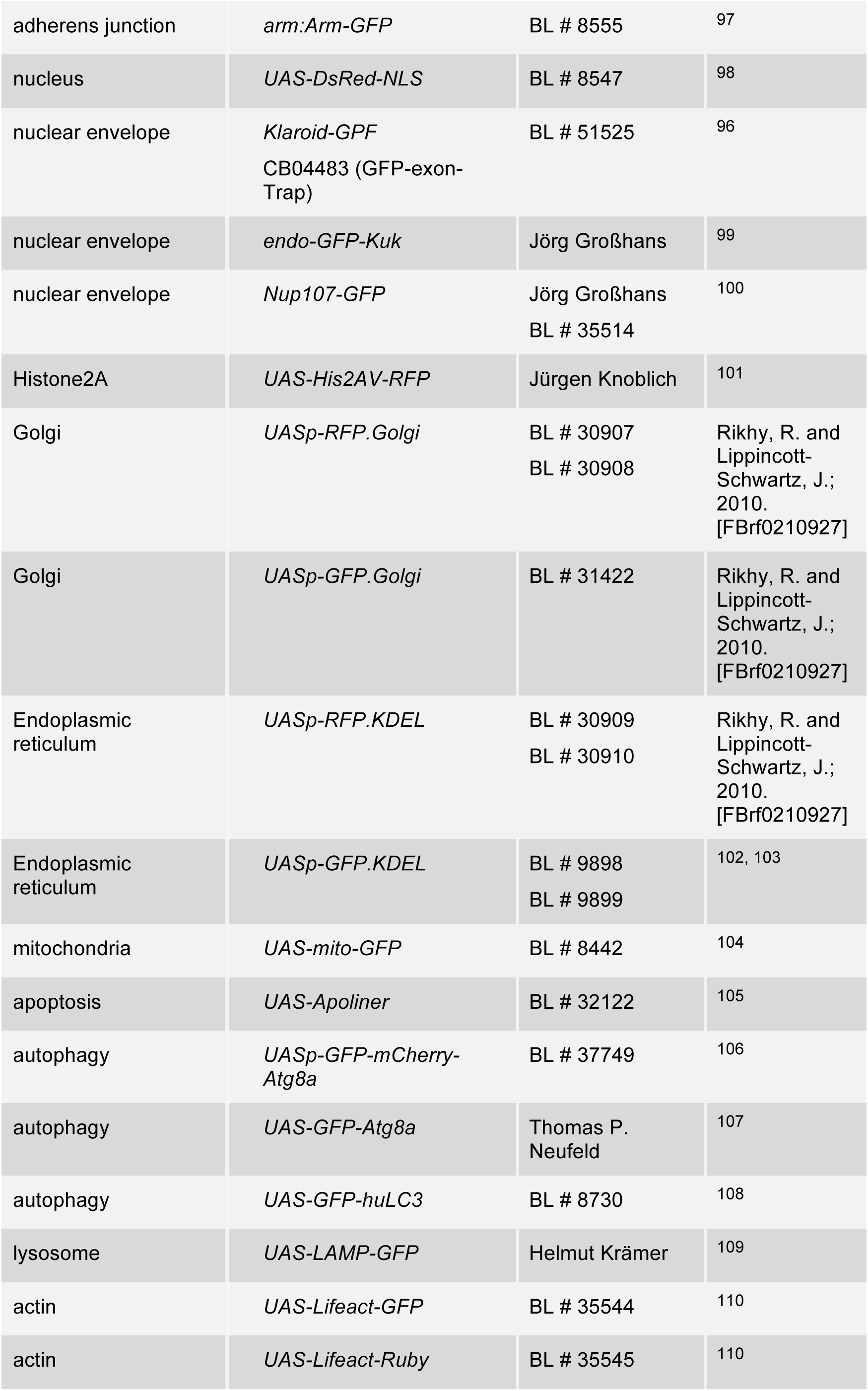
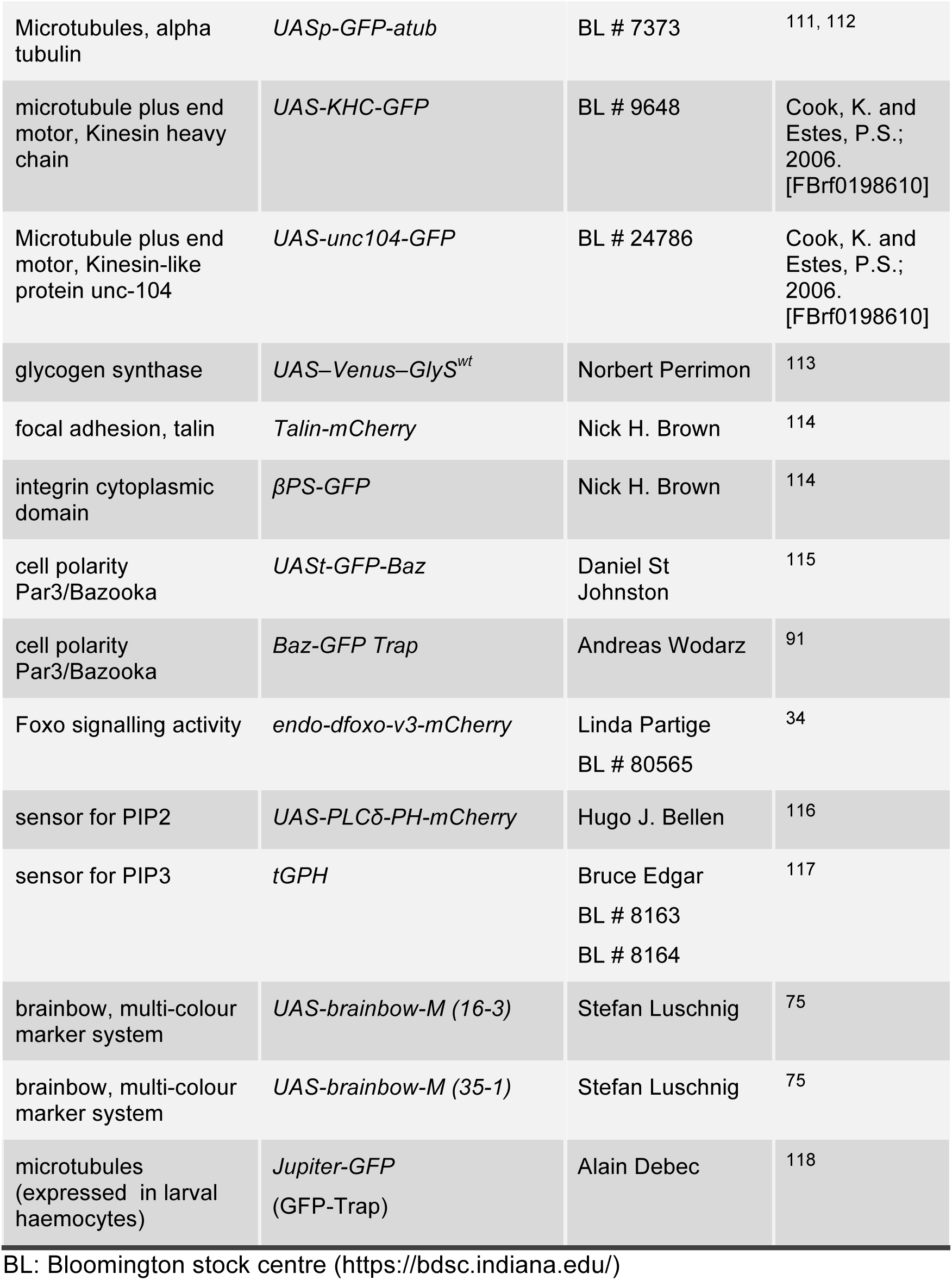
Fly stocks for imaging subcellular compartments in Drosophila larvae.

### Microscope

- For all live imaging we used an inverted spinning disk confocal microscope (Inverse, Nikon TiE, Yokogawa CSU-X1) with the Nanopositioning Piezo Z stage control system (NanoScan OP400, Prior Scientific), Plan-Fluor 40x/1.3 NA Oil-immersion DIC objective or Plan-Apochromat 60×/1.2 NA water-immersion objective; EMCCD camera (C9100-50 CamLink; 1000 × 1000 pixel) controlled by the Volocity software 6.3.57, 488 nm and 561 nm channels with appropriate bandpass filters and dichroic mirrors.
- The microscope setting was optimized to perform the long-term 3D live imaging (1-8 hours, with 2-15 min time interval) or fast imaging (with a time interval of some ms) outlined in this protocol as described in the **Procedure** section.

### Ultraviolet laser

- All laser ablation and wounding experiments were performed with 60×/1.2 (for single-cell ablation) or 40x/1.3 (for multi-cell ablation) objective lenses on a spinning disk microscope, which was equipped with 355 nm pulsed ultraviolet laser (DPSL-355/14; Rapp Opto-Electronic, 14-mW mean power, 70-µJ per pulse). Before starting the laser wounding experiments, the laser was calibrated. Parameters used to induce laser-wound or laser-cut on cell-cell junction are described in the **Procedure** section.

### Image-processing and quantification software

- Volocity software 6.3.57 from PerkinElmer (https://www.perkinelmer.de/lab-products-and-services/resources/whats-new-volocity-6-3.html).
- ImageJ or Fiji, National Institutes of Health (https://fiji.sc/ or https://imagej.net/Fiji).

### Anaesthetics

- Diethyl ether, for analysis Merck Cat. No.: 1.00921.1000 (http://www.merckmillipore.com/DE/de/product/Diethyl-ether,MDA_CHEM-100921?ReferrerURL=https%3A%2F%2Fwww.google.com%2F)
- Desflurane Baxter (Suprane® Inhalation Anaesthetic Desflurane Inhalation Liquid Bottle 240 mL; NDC: 10019064134) (https://mms.mckesson.com/product/842158/Baxter-10019064134)
- Isoflurane Baxter (FORANE (isoflurane, USP) Liquid for Inhalation 250 mL Colored Glass Bottles; NDC: 10019-360-60) (https://ecatalog.baxter.com/ecatalog/loadproduct.html?cid=20016&lid=10001&hid=20001&pid=822726)
- Ethyl acetate, Sigma-Aldrich; EC Number 205-500-4 (https://www.sigmaaldrich.com/catalog/product/mm/100864?lang=de&region=DE&gclid=EAIaIQobChMIsdbhqvN4gIVj5AYCh2KZADAEAAYASAAEgLdfPD_BwE)

### Equipment

The equipment is shown in **Fig. 2**. The numbers in the list correspond to the magenta numbers in **Fig. 2**.

1. 10 ml plastic pipette (polyethylene): (https://oman.desertcart.com/products/39917106-gle2016-clear-white-plastic-liquid-dropper-pasteur-disposable-graduated-transfer-pipettes-pipetting-10ml-100-pack)
2. Diethyl ether, Merck Cat. No. 1.00921.1000:
3. (http://www.merckmillipore.com/DE/de/product/Diethyl-ether,MDA_CHEM-100921?ReferrerURL=https%3A%2F%2Fwww.google.com%2F)
4. Glass Wheaton Coplin staining jar with PP screw cap and 55 ml or 60 ml capacity: (https://www.capitolscientific.com/Wheaton-Science-900570-Coplin-Staining-Jar-with-PP-Screw-Cap-60mL-Capacity-Holds-5-to-10-Slides)
5. Plastic Petri dish, diameter 100 mm, polystyrene, P5731 Sigma: (https://www.sigmaaldrich.com/catalog/product/sigma/p5731?lang=de&region=DE&cm_sp=Insite-_-prodRecCold_xviews-_-prodRecCold3-1)
6. Plastic Petri dish, diameter 60 mm, polystyrene, P5481 Sigma: (https://www.sigmaaldrich.com/catalog/product/sigma/p5481?lang=de&region=DE)
7. Culture dish for imaging, CELLview cell culture dish, PS, 35/10 mm glass bottom, 1 compartment, sterile (Greiner Bio-One Item No: 627861): (https://shop.gbo.com/de/germany/products/bioscience/zellkultur-produkte/cellstar-zellkultur-schalen/cellview-zellkultur-schale-mit-glasboden/627861.html)
8. Culture dish with 4 compartments, ideal for imaging of 2 to 4 different genotypes at the same time. CELLview cell culture dish, PS, 35/10 mm glass bottom, 4 compartments, TC, sterile (Greiner Bio-One Item No: 627870): (https://shop.gbo.com/de/germany/products/bioscience/zellkultur-produkte/cellstar-zellkultur-schalen/cellview-zellkultur-schale-mit-glasboden/627870.html)
9. Tap water at room temperature
10. Empty food vial (to raise the larva after imaging), one for each larva
11. 10 ml glass beaker (Duran, Cat. No. 21 106 08): (http://www.duran-group.com/en/products-solutions/laboratory-glassware/products/boiling-flasks-and-general-laboratory-glassware/beaker.html)
12. Black nylon sheet, for better visualization of larvae during washing and sorting, (black plastic file folder): (https://www.viking.de/de/p/4191-SZ?cm_mmc=Google-_-PLA_GEN_GOOGLE-SHOPPING_ordnung-im-buero_-_-ordnung+im+buero-_-PRODUCT_GROUP&&cm_mmc=Google-_-PLA_GEN_GOOGLE-SHOPPING_ordnung-im-buero_-_-ordnung+im+buero-_-4191-SZ&gclid=EAIaIQobChMI68r8obKY4wIVhIbVCh2uAQauEAQYCiABEgIH6PD_BwE&gclsrc=aw.ds)
13. Self-made larval cage with lid (Fig. 2, 3)
14. Paper napkin (with little fluff)
15. Timer
16. Two paint brushes with real hair (not plastic): one for washing the larvae (size 1; Cat. No. 8-22621) and the finer one for sorting and orienting them (size 2/0; Cat. No. 8-69656): (https://www.gerstaecker.de/da-Vinci-Serie-1520-Rotmarder-Aquarellpinsel.html)
17. Spatula
18. Cotton wool
19. Filter forceps, for transfer from larval cage into and out of the anaesthetic chamber
20. Nylon mesh, 300-400 µm pore size. e.g Nylon mesh 400µM “Nybolt polyamid MONO NO: PA 400/47 or Nylon filtration tissue (sifting fabric) NITEX, mesh opening 300 µm (Biolab, ID: 074003)
21. 1.5 ml microcentrifuge (Eppendorf Safe-Lock Tubes)
22. Bunsen burner

## Procedure

### Selection and generation of optimal lines for live imaging TIMING variable

1 To image epidermal wound healing in larvae, use either: *w^1118^; e22c-Gal4, UAS-Src-GFP, UAS-DsRed2-Nuc/CyO; +* or: *w^1118^; +; A58-Gal4, UAS-Src-GFP, UAS-DsRed2-Nuc/TM6B* flies. In both lines, the plasma membrane is marked by Src-GFP and nuclei by DsRed2-Nuc^34, 61^. Both drivers, *e22c-Gal4* and *A58-Gal4,* are expressed in the epidermis. *e22c-Gal4* is expressed from embryonic stage 10-11 onward and *A58-Gal4* from the first larval instar onward^61, 70^.
2 To visualise the adherens junctions for laser ablations at cell interfaces, use a transgenic line, in which endogenous DE-Cad is marked by GFP (*w^1118^, y^-^; endo:DEcad-GFP; +*)^71^. We obtained best results with lines in which DE-Cad is expressed under its own promoter rather than a tubulin or ubiquitin promoter. The *endo:DE-Cad-GFP* chromosome is homozygous viable and causes no cellular or developmental abnormalities. The transgenic proteins are sufficiently bright to be imaged easily even when the animal is heterozygous for the chromosome.

### Generation of larvae for wound healing and laser cutting TIMING variable (2-3 days)

3 **Wound healing:** Both *e22c-Gal4, UAS-Src-GFP, UAS-DsRed2-Nuc* and *A58-Gal4, UAS-Src-GFP, UAS-DsRed2-Nuc* chromosomes are kept over a balancer. For imaging of the control animals cross them with wild type (*w^1118^; +; +*):
**P:** virgin female *w^1118^; e22c-Gal4, UAS-Src-GFP, UAS-DsRed2-Nuc/CyO; +* x **P:** male *w^1118^; +; +* **F1:** w^1118^; e22c-Gal4, UAS-Src-GFP, UAS-DsRed2-Nuc/+; + Or **P:** virgin female *w^1118^; +; A58-Gal4, UAS-Src-GFP, UAS-DsRed2-Nuc/TM6B* x **P:** male *w^1118^; +; +* **F1:** w^1118^; +; A58-Gal4, UAS-Src-GFP, UAS-DsRed2-Nuc/+ The imaging is performed with the larvae from the F1 generation.
4 **Laser cut:** For laser ablations in which cell outlines are labelled by *endo:DE-Cad-GFP*, set up the following cross with wild type (*w^1118^; +; +*): **P:** virgin female *w^1118^, y^-^; endo:DEcad-GFP; +* x male *w^1118^; +; +* **F1:** w^1118^; endo:DE-Cad-GFP/+; +

### Developmental synchronization of the larvae TIMING 5-6days

5 To obtain a sufficient number of animals at the proper developmental stage (early or mid-L3 instar larval stage) the larval growth has to be synchronised. One day after setting up a cross, let the flies lay eggs in food vials for 30-60 min at 25°C and repeat this several times and in parallel in several vials and keep them at 25°C. Then collect the L3 larvae 5.5 to 6 days after egg deposition. Not all stocks develop at precisely the same speed, especially if they carry mutations or complex combinations of chromosomes. Thus, this timing has to be carefully determined for every stock. **ALTERNATIVE** A more stringent method for an exact synchronisation is following the protocol of Maimon, et al.^74^: depending on the size of the fly vial, cross 20-30 males with 60-80 virgin females (2-7 days old) in a fly bottle and keep for 3-4 days at 25°C. For exact synchronisation of larvae let the flies lay eggs for 2-4 hours and remove the flies after that. Collect 60-80 1^st^ larval instar stage 24 h after removal of the flies (i.e. approximately 24h after egg laying (AEL) with a two-hour interval for egg deposition), transfer to new food vials and place in an incubator at 25 °C. Collect 25 to 30 larvae between 72-85 h AEL for live imaging.
6 All fly stocks, crosses and larvae have to be maintained at 25 °C under a 12:12 h light/dark cycle at constant 65% humidity on standard fly food.

### Construction of the larval cage and anaesthetic chamber TIMING ∼15 min

**Construction of larval cage with lid (Fig. 3):**

7 To make a larval cage (Fig. 3), heat the scalpel using a Bunsen burner (Fig. 3a) and cut a 1.5 ml microcentrifuge tube 5 mm from the top (Fig. 3b). Melt carefully the cut surface of the top of the tube with the cap and glue it immediately to a nylon mesh with 300-400 µm pore size (Fig. 3c). Wait 5 min and then cut off the overhanging parts of the nylon mesh with a scissor (Fig. 3d).

### Construction of anaesthetic chamber (Fig. 4)

8 To make an anaesthetic chamber (Fig. 4c) plug tightly the glass Wheaton Coplin jar with cotton wool (Fig. 4a). To minimize the amount of anaesthetic to be used, the cotton wool should be pressed very tightly. The maximum volume possible in the Coplin jar should be occupied by compacted cotton wool, (Fig. 4a-b) leaving only enough space for a 10ml glass beaker (Fig. 4f).
9 Plug also the 10 ml glass beaker with cotton but keep enough space to place the larval cage into it (Fig. 4b). From this step onwards work under the fume hood. **! CAUTION** Exposure to anaesthetic gas is a serious safety concern. The diethyl ether or any other anaesthetic must be directly vented out of the room using a certified ventilated fume hood or a bio-safety cabinet. **! CAUTION** Diethyl ether is sensitive to light and inflammable. It can form an explosive vapour and it should kept dark and cold under a ventilated fume hood. This is also the case for all other anaesthetics we have listed in this protocol: desflurane, isoflurane, ethyl acetate.
10 Fill the Coplin jar with the help of a large 10 ml plastic pipette (polyethylen) with 15-20 ml diethyl ether (Fig. 4d). **Δ CRITICAL STEP** The amount of diethyl ether required depends on the free volume remaining in Coplin jar after plugging with cotton wool and may vary (15-20 ml). There should be enough liquid diethyl ether inside the chamber to saturate the chamber with diethyl ether vapour.
11 Place the glass beaker into the Coplin jar (Fig. 4e).
12 Fill also the glass beaker with enough diethyl ether to only just soak the cotton wool (in our experience, 2-3 ml; Fig. 4f). **Δ CRITICAL STEP** The cotton wool inside the glass beaker should only be soaked, there should be no free liquid. **! CAUTION** The diethyl ether inside the Coplin jar should not rise above the middle of the glass beaker, to prevent either flowing into the glass beaker and into the larval cage.
13 Close the Coplin jar tightly. **! CAUTION** Diethyl ether and all other anaesthetics listed in this protocol rapidly evaporate, so that in each step the anaesthetic chamber should be opened for the absolute minimum of time, and immediately shut again. **! CAUTION** The anaesthetic chamber (Fig. 4c) always has to be closed tightly with its cap and kept under the fume hood, also when no experiments are performed. **! CAUTION** Diethyl ether is a solvent and can dissolve some synthetic materials. Therefore its direct contact with plastic (e.g. polystyrene) should be avoided. **Δ CRITICAL STEP** Always keep the anaesthetic chamber and larval cage in a dry environment and avoid any contact with water. The diethyl ether does not anaesthetise reliably if the larvae are damp or wet. The dry air under the fume hood mitigates against this. **ALTERNATIVE** For long-term immobilisation we used diethyl ether as anaesthetic. For short- and mid-term (<60min) live imaging the same anaesthetic chamber and larval cage can be used with any of the other anaesthetics (Table 1, 2 and Fig. 6). **PAUSE POINT**

### Anaesthetisation and mounting the larvae for imaging TIMING 8-15 min

14 To wash the larvae place both big (100 mmm) and small (60 mm) petri dishes on top of a black nylon sheet (Fig. 5a). The black nylon sheet helps to see and sort the larvae. Fill both petri dishes with tap water (room temperature). To reduce any chlorine in the tap water, we recommend keeping the water for some days at room temperature, so that the chlorine evaporates.
15 Scoop the L3 instar larvae with a metal spatula out of the soft food into the big petri dish (Fig. 5a) and wash them thoroughly by gently moving them in the water with the paint brushes (size number 1) to remove the food from the larvae (Fig. 5b).
16 Transfer the larvae with the same paint brush into the smaller petri dish to remove the leftover food also by gently moving them in water (Fig. 5b-c).
17 Collect 20-30 larvae one by one with the same paint brush and dry them on the paper napkin (Fig. 5c). **Δ CRITICAL STEP** This step is essential and could affect the anaesthetisation. Since diethyl ether and all other anaesthetics we have tested here are hydrophobic, the larvae must be gently but thoroughly dried. The moisture of the environment, including that of the larval cage, or larva itself prevents the diethyl ether to enter through the outer cuticle or the tracheal system and to become effective. **? TROUBLESHOOTING** **Δ CRITICAL STEP** In order to limit any possible stress, keep the washing and drying steps as short as possible (< 5 min).
18 Transfer the washed and dried larvae into the cap of the larval cage (Fig. 5d). If necessary, also in this step dry the larvae and the cage carefully with a paper napkin.
19 To expose the larvae to diethyl ether, open the anaesthetic chamber; place the larval cage with the cap facing down (with the larvae inside the cap) inside the glass beaker using the filter forceps (Fig. 5e). **Δ CRITICAL STEP** Close the cap of the larval cage tightly to avoid the contact of the larvae with the diethyl ether-soaked cotton wool inside the glass beaker. Contact with diethyl ether can induce a cold shock in larvae, induce stress or even death.
20 Close the screw cap of the anaesthetic chamber tightly (Fig. 5f). Late L2 and early L3 larvae can be exposed to diethyl ether for 3 or 3.5 min respectively. Middle or late L3 larvae can be exposed to diethyl ether for 4-4.5 min (or at most 5min) (Table 2). However, these times may have to be adjusted for genetically modified or mutant larvae, especially if the genetic condition affects body size. **ALTERNATIVE** The chamber can also be used for desflurane, ethyl acetate or isoflurane. Exposure times for these anaesthetics vary and also depend on larval growth (size and larval stage). For the exact exposure time see **table 2**. **! CAUTION** The time of exposure to diethyl ether negatively correlates with survival rates of the larvae. To minimize the lethality the exposure time has to be restricted to the exact time mentioned above and in Table 2. **? TROUBLESHOOTING**
21 After the indicated time, open the anaesthetic chamber and remove the larval cage with filter forceps.
22 Transfer the anaesthetised larvae one by one by using the moistened fine paintbrush (size 2/0) into a culture dish for imaging with the 35/10 mm glass bottom (Fig. 5g).
23 With the same fine paintbrush sort and orient the larvae under the stereomicroscope (Fig 5g-h). If needed, the larvae can be also sorted for fluorescent markers under a fluorescence stereomicroscope before imaging. If larvae stick to each other and are not easy to move and orient, moisten the brush with tap water. However, after all larvae are in the right position, remove the leftover water carefully with a paper napkin. **Δ CRITICAL STEP** The larvae should be placed at a distance of 2-3 mm to each other (Fig. 5h). As soon as one larva starts to move or is not completely anaesthetised, it should be removed from the culture dish, otherwise it wakes up the other larvae. If one larva wakes up, some of the other larvae also wake up, so that it is important to remove any moving larvae immediately to avoid a domino effect. This holds also during imaging. The effect is not caused by the larva moving and touching other larvae, but by an unknown mechanism that does not require direct contact. **? TROUBLESHOOTING**
24 To induce a laser wound in the dorsal epidermis and record the healing process with an inverted microscope, face the dorsal side of the larvae to the glass bottom of the culture dish if using an inverted microscope (as we did; Fig. 5h). To know the correct orientation of the recorded time lapse always orient the larvae in the same direction for all experiments; for instance, with the head always pointing to the right side (Fig. 5h-i). **ALTERNATIVE** For imaging of different organs, larvae have to be oriented and imaged from the appropriate side. Image the (i) epidermis, neurons, muscle, from the dorsal side, (ii) midgut, wing imaginal disc, peripheral neurons, lateral pentascolopidial (Lch5) organ, from the lateral side, (iii) brain and ventral nerve cord from the ventral side. Trachea, fat body and haemocytes can be imaged from any orientation. **! CAUTION** An indication whether the anaesthetised larvae are alive is their heartbeat. The heartbeat can easily be seen from the ventral or lateral side of the larva under the stereomicroscope. Larvae without heartbeat should be removed from the experiment.
25 With an inverted microscope there is no need to use mounting media, oil or coverslip. The cuticle of the larvae is made of chitin, a long linear sugar^73^. As soon as larvae are dry and are not in contact with water or in a humid environment, they stick to the glass. Therefore, there is no need to use adhesive or glues to stick them to the culture dish (Fig. 5h-i). **Δ CRITICAL STEP** In addition to the fact that water prevents adhesion of the cuticle to the glass it could also wake up the larvae. Contact with water should be avoided until the end of the experiment. **ALTERNATIVE** If using an upright microscope there is no need to mount the larvae with mounting media. Put a coverslip on top of the larvae, which should have the thickness suited for the objective lens. Avoid pressing larvae with the coverslip.
26 Now the samples are ready for imaging (Fig. 5i).

### Microscope parameters used for optimal resolution and live confocal imaging TIMING variable (up to 8 hours)

27 For all our live imaging we used an inverted spinning disk confocal microscope as described above (section **Materials/Microscope**)^34, 35^.
28 Microscope setting for live imaging of *A58-Gal4, UAS-Src-GFP, UAS-DsRed2-Nuc* or *e22c-Gal4, UAS-Src-GFP, UAS-DsRed2-Nuc* larvae is: 488 nm and 561 nm channels (exposure time 100-180 ms for both 488 nm and 561nm laser beam); 55-65 z-stacks with a step size of 0.28 µm with 60x/1.2 and 0.24 µm with 40x/1.3 (covering 15-20 µm depth) were taken every 2– 15 min for a 1–8h. This setting does not induce phototoxicity or strong photobleaching during 8 hours of live imaging. In our experiments, we used the 488 nm laser at 12–15% power, and the 561 nm at 2–4%. However, since every laser is different, and lasers change during their life, and because different constructs have different fluorescent properties, these values have to be individually determined. **Δ CRITICAL STEP** To protect the samples against phototoxicity and photobleaching during imaging, reduce the laser power and increase the exposure time (100-200 ms). Longer exposure time obviously reduces the speed of imaging and is not suitable for imaging with narrow time intervals (less than 1min).
29 Microscope setting for live imaging and laser ablation of *endo:DE-Cad-GFP,* larvae is: 488 nm (10-30% laser power in our case) channel (exposure time 15-60 ms); single planes were taken every 0.5 s for 5–7 min, starting ∼2 min before ablation, and finishing ∼5 min after ablation.
30 The settings in **step 29** are also suitable for monitoring fast cellular and subcellular events, e.g. vesicle trafficking or calcium flashes, which require high-speed imaging at narrow time intervals (some ms). **Δ CRITICAL STEP** In general, reducing the exposure time and minimizing the number of z-stacks will allow recording at shorter time intervals. However, for better signal to noise ratios, higher laser power is desirable. **! CAUTION** Increasing the power of the laser beam increases photobleaching and phototoxicity and is only recommended for short-term live imaging. It is important for each set of experiments with different fluorescent markers first to run test experiments to find the optimal parameters.

### Live imaging of internal organs TIMING variable (5-120 min)

31 For imaging internal organs the spinning disk microscope with the settings described in **step 29** is recommended.
32 For optimal tracking of the organ of interest reduce the number of z-stacks or choose single plane imaging. Combine this with a short exposure time (15-60 ms).
33 If necessary, the imaging of different planes can be carried out by manually changing the fine-focus during live imaging (Video 3, 7, 8, 11). This depends on the ability of the software to record and transfer the images simultaneously. This is possible for the Volocity software 6.3.57 we have used.
34 For analysis and quantification it is important to have the same experimental conditions for all genotypes and also the same image settings.

### Multi- and single-cell laser ablation for wound healing studies TIMING 1sec-10min

35 To image events immediately after ablation use the same microscope and objective lens for laser ablation and imaging.
36 Perform the laser ablation with a 60×/1.2 (for single-cell ablation) or 40x/1.3 (for multi-cell ablation) objective lens on the spinning disk microscope equipped with a 355 nm pulsed ultraviolet laser (DPSL-355/14; Rapp Opto-Electronic, 14-mW mean power, 70-µJ per pulse).
37 Before starting the experiment, the laser should be correctly calibrated. For a high laser power efficiency, the microscope focus during calibration is extremely important and should be adjusted on the plane nearest to the glass of imaging dish, e.g. on the apical region of the epidermis to be wounded.
38 To start the laser ablation, turn on the UV laser and all associated devices. Use an appropriate dichroic mirror to reflect the UV laser into the specimen. To be on safe side change the shutter to allow the bypass of UV laser. **! CAUTION** Make sure to avoid exposure of your eyes or other body parts to the UV laser beam through the objective lens or ocular. The reflection of light from the laser beam can cause serious damage to the eyes or skin.
39 Set all laser parameters.
40 Find the dorsal midline of abdominal segment A3 or A4 (or any other region of interest),
41 Focus on the epidermal cells and make a single image to record the pre-wounding situation. Pause the image acquisition.
42 To induce a single-cell ablation, focus on the nucleus in the target cell and mark three spots (diameter for each spot is 6 px) on the nucleus with the software of the laser (Fig. 10, Supplementary Video 12a). **ALTERNATIVE** The spot size of the UV laser beam can vary between 1 to 10 px depending on cell size. For cells of 20-30 µm diameter adjust the UV spot to 1-2 px diameter and for bigger cells (40-80 µm diameter) to 3-6 px diameter.
43 To deliver a laser pulse, click the appropriate icon to release the laser power and shoot the cells with the UV laser beam with 1 pulse/µm laser power of ∼0.30 µJ energy. The pulse energy is measured on the objective lens using a half wave plate and a polarizer form 50 nJ to 12 µJ. This should generate a damaged area of 40-60 µm diameter (800-2500 µm^2^ area). **ALTERNATIVE** We performed laser ablation during time-lapse imaging without pausing the image acquisition. Alternatively, one can also pause imaging, perform the laser ablation and then continue imaging. **Δ CRITICAL STEP** The highest power of laser beam efficiency is on the focal plane where the beam was calibrated. Therefore, to ablate the cell(s), the correct focal plane has to be selected. If laser wounding is unsuccessful, first change the focal plane by changing the fine focus and repeat the laser treatment. If this does not help, gradually increase the laser power and try again until you see an effect. Alternatively, calibrate the laser again or contact the provider to help you for the correct calibration, because this is not trivial. **? TROUBLESHOOTING** **Δ CRITICAL STEP** If the laser target site is too close to a neighbouring cell, this can result in damage of the cell membrane, with deleterious effects on and possibly death of the neighbouring cells. If the target site in the cell to be killed is not at the correct level or position, and only cytoplasm is damaged, rather than the nucleus, this may not reliably lead to cell death; sometimes the cells are able to heal themselves, in a process often associated with accumulation and contraction of actomyosin. In contrast, laser ablation of the nucleus always immediately leads to cell death and a response from the neighbouring cells, even without damage to the neighbouring cells.
44 For multi-cell ablation, one laser pulse on a spot of 6 px diameter was delivered to the nucleus of each cell to be killed and to each of their common junctions (Fig. 10, Supplementary Video 12b). A multi-cell laser ablation (3-8 cells) can generate a wound of 80-120 µm diameter (3000-8500 µm^2^ area). **ALTERNATIVE** With a spinning disk wide confocal microscope larger wounds can be induced, because the field of view is 4 times larger than a standard scanning head. **! CAUTION** The pulse and power of the UV laser beam is the same for both single- and multi-cell ablation (1 pulse/µm, ∼0.30 µJ, energy measured on the objective lens). The only difference is the number of positions shot by the laser. For single-cell ablation, 3 partially overlapping spots and for a multi-cell ablation, e.g. ablation of three cells, 7 different spots have to be marked to induce a wound. In all cases, the diameter of the spots was 6 px.

### Laser ablation of the cell boundary to measure junctional tension TIMING 1sec-10min

45 To perform a laser cut at the plane of the cell-cell junction and to avoid a wound healing response use the following UV laser beam setting: 1 pulse/µm of ∼0.25 µJ energy (measured on the objective lens).
46 Find the dorsal midline of abdominal segment A3, A4, or A5 the microscope.
47 Focus on the most apical side of epidermal cells.
48 Mark the cell boundary with a single spot (diameter for each spot size is 1 px).
49 The recoil occurs within some ms, therefore the laser cut has to be induced during the time-lapse imaging, to capture and quantify the immediate response to cutting. To image as fast as possible set up the microscope for single plane and image only one channel (488 nm, green for *endo:DE-cad-GFP*).
50 Before UV laser ablation, first record a time lapse of ∼2 min (time interval every 0.5 sec).
51 To perform the laser cut, press the appropriate icon to release the laser power (1 pulse/µm of ∼0.25 µJ energy, measured on the objective lens). **Δ CRITICAL STEP** If the laser cut is unsuccessful, change the focal plane or increase the laser power or both.

### Monitoring the survival of the animals and quantitative evaluation of data TIMING 2-5 days

52 To monitor viability after imaging, transfer each larva individually to an empty fly food vial.
53 Record on the vial the same name/ID the larva that was used for recording the imaging data.
54 Allow the larva to recover and develop under normal conditions: at 25 °C under a 12:12 h light/dark cycle at constant 65% humidity.
55 Analyse the imaging data only from those animals that survive to pupa or adult flies. This helps to distinguish the effects of the experimental variables (through genetic manipulation or drug treatment) from the overall fitness of the animals and to avoid analysing artefacts.

### Import and processing of the imaging data TIMING variable

56 For converting, processing and importing data any suitable software can be used: Volocity, Imaris, or Fiji. We use Volocity software (6.3.57 version).
57 The imaging parameters described in this protocol are adjusted to each fluorescent dye and optimized for long-term live imaging, so that fluorescent signal intensity is maintained until the end of live imaging (8 h) and is suitable for qualitative and quantitative analysis.
58 The quality of images taken with the microscope settings described here are in general good enough for quantification and analysis.
59 If necessary, the threshold of fluorescence signals can be post-processed in a linear fashion.
60 For analysis and quantification of the data by Fiji or Imaris, the data have to be converted to “ome.tiff” before import to those programs.

### Analysis and quantification of wound closure and junctional tension TIMING variable

61 For 4D data analysis the quantification of images, use Volocity software (6.3.57 version), Fiji (Fiji Is Just ImageJ; National Institutes of Health) or Imaris (Bitplane)
62 For correct measurement, import the metadata to the software (Volocity, Fiji, or Imaris), where the data can be analysed.

#### Wound healing dynamics

63 Wound areas, defined as the area left open by the dead cells between the lamellipodia. Measure this area manually using Volocty, Fiji or Imaris^34^.

#### Cell tension

64 Measure the displacement of the cell vertices after laser ablation of the cell bounds over a time interval of 2-5 sec, where the first measurement point is the time of ablation^35^.

## Timing

Steps 1-4, selection and generation of optimal lines for live imaging: variable 2-3 days.
Steps 5-6, staging and synchronizing larval growth: 5-6 days.
Steps 7-13, construction of larval cage and anaesthetic chamber: 15 min.
Steps 14–26, anaesthetisation and mounting of larvae: 8-15 min.
Steps 27–34, setting up the microscope and long-term imaging of epidermis or internal organs: variable up to 8 hours.
Step 35-44, multi- and single-cell laser ablation for wound healing studies: 1 sec-10 min.
Step 45-51, laser cutting of cell boundaries to measure junctional tension: 1 sec-10 min.
Steps 52-55, monitoring the survival of the animals and evaluation of data for quantitative measurements: 2-5 days
Steps 56-60, import and processing of images: variable.
Step 61-64, analysis and quantification of wound closure and junctional tension: variable.

## Troubleshooting

Troubleshooting advice can be found in **Table 5**.

**Table 5.** Troubleshooting table.

| step | problem | possible reason | solution |
| --- | --- | --- | --- |
| 17 | Larvae are not immobile after anaesthetisation. | i) Larvae or equipment, in direct contact with larvae, are not completely dry<br>ii) Larvae are too crowded.<br>iii) Exposure to anaesthetic is too short | i) Dry the larvae, larval cage, paintbrushes and imaging dish properly with a paper napkin. Make sure the microscope chamber does not produce any humidity, which can occur especially if the equipment was previously used with *ex vivo* or *in vitro* materials.<br>ii) Avoid putting more than 30 larvae (late L2 or early L3) or 20 (middle L3) larvae in a larval cage for anaesthetisation.<br>iii) Observe the exact times mentioned in table 2. |
| 20 | Larvae are not vital; death rate is high during imaging. | Larvae were exposed too long to the anaesthetic | Observe the exact times mentioned in table 2.<br>Reduce the exposure time to the anaesthetic if using weak mutant lines. |
| 23 | Larvae start moving during imaging. | Larvae in the imaging dish are too crowded | Place the larvae at a distance of 2-3 mm to each other and not less than that |
| 43 | UV laser wounding is unsuccessful. | The calibrated focal plane is different than the focal plane of imaged larvae.<br>The device is not correctly calibrated. | Change the fine focus on focal plane.<br>Calibrate the laser again or contact the provider to help you for correct calibration. |

## Anticipated results

Live imaging of systemic, cellular and subcellular processes with high spatio-temporal resolution can improve our understanding of those processes. This protocol allows long-term live imaging of larval organs, subcellular organelles and events in *Drosophila* larvae (Fig. 7, Fig. 10 and Supplementary Video 1-12).

**Fig. 10.**
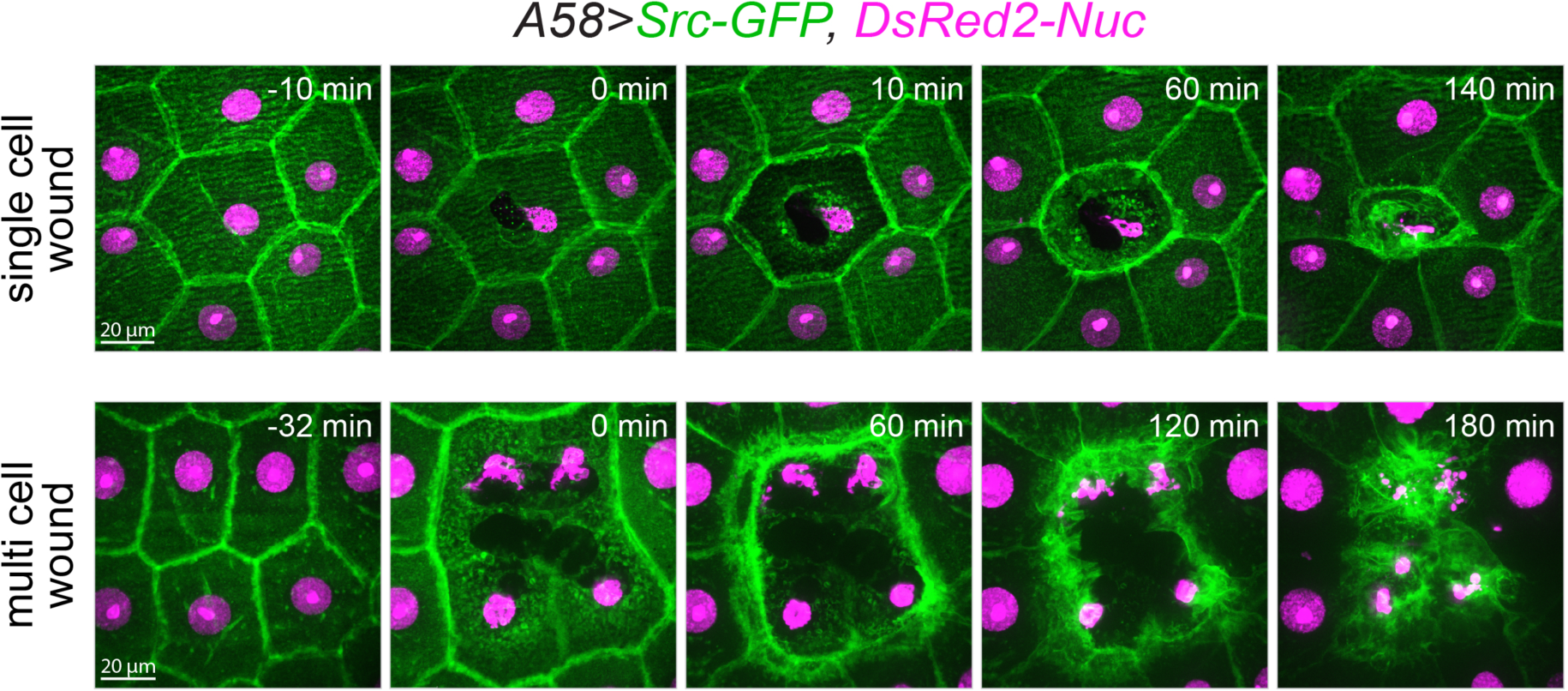
Single- and multi-cell laser wounding in the larval epidermis. Time-lapse series of single-cell and multi-cell wound healing in L3 larvae expressing Src-GFP (green) and DsRed2-Nuc (magenta) to mark the cell membrane and nuclei in the epidermis: *A58>Src-GFP; DsRed2-Nuc (w^1118^; +; A58-Gal4,UAS-Src-GFP, DsRed2-Nuc/+*). Projections of a time-lapse series in early L3 larvae. Each frame is a merge of 57 planes spaced 0.28 µm apart. Scale bars: 20 µm. Corresponds to **supplementary Video 12.**

The method has enabled us to detect novel spatio-temporal details of insulin and TOR signalling and their consequences in the wound healing process^34^. We observed activation of insulin receptor signalling (PI3K/FOXO) within minutes in the epidermis (Fig. 11a, Supplementary Video 13). The epidermis-specific interference with Insulin receptor signalling reduced PI3K signalling (Fig. 11b, Supplementary Video 14) and caused a delay of epidermal repair^34^. We found that the insulin signalling network is needed for the efficient assembly of an actomyosin cable around the wound, and constitutively active myosin II regulatory light chain suppresses the effects of reduced IIS^34^.

**Fig. 11.**
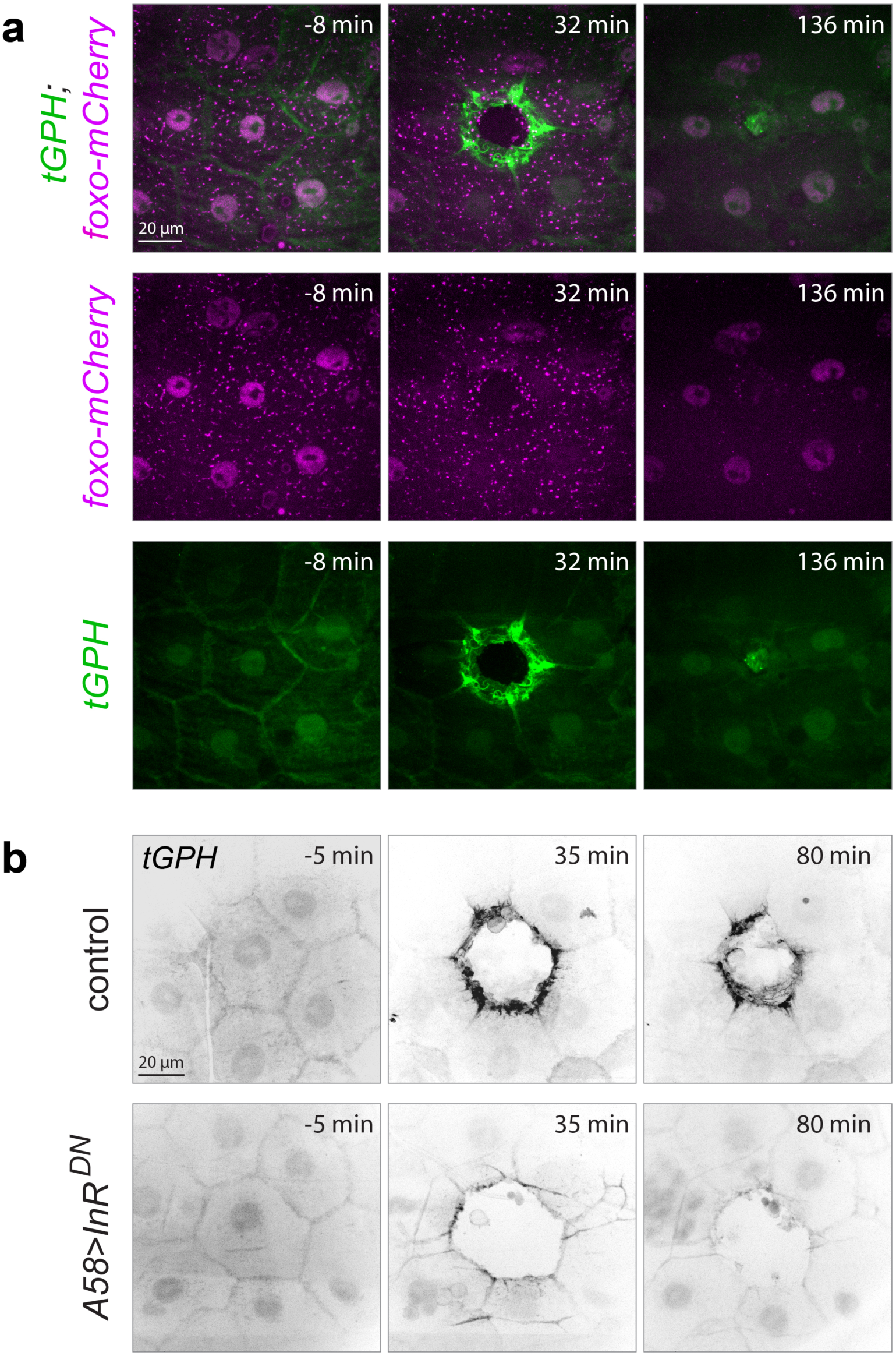
Live imaging of insulin/PIP3/FOXO signalling and the effect of reduced insulin receptor signalling on PIP3. (**a**) Time lapse of a single-cell wound in larva expressing tGPH (*tubulin:GFP-PH*) (green), a PIP3 reporter and endogenous FOXO-mCherry (magenta). PIP3 accumulation (green) and dynamic FOXO (magenta) shuttling between nucleus and the cytoplasm in cells directly surrounding the epidermal wound of L3 larvae during wound healing. (**b**) Distribution of tGPH (black) in the epidermis of control and *A58>InR^DN^* larvae. In *A58>InR^DN^* larvae a dominant negative version of the insulin receptor (InR^DN^) is expressed under the control of the *A58-Gal4* driver. Reduced insulin receptor signalling (*A58> InR^DN^*) in the epidermis of larvae lowered the PIP3 accumulation (grey) at the wound edges and led to wound healing delay. (**a-b**) Projections of a time-lapse series of the wounded epidermis in early third instar larvae. Each frame is a merge of 57 planes spaced 0.28 µm apart. Scale bars: (**a,b**) 20 µm. Transgene genotypes: (**a**) *tGPH; foxo-mCherry* (*w^-^; tGPH; endo-dfoxo-v3-mCherry*) and (**b**) control (*w^1118^; tGPH/+; A58-Gal4/+*) and *A58>InR^DN^*(*w^1118^; tGPH/+; A58-Gal4/UAS-InR^DN^*). Corresponds to **supplementary Video 13 and 14.**

In summary, the immobilisation method and long-term *in vivo* imaging described here, coupled with genetic manipulations allows the study of many aspects of biology and physiology.

## Supporting information

Supplementary Video 1 | Live imaging of mitochondria during epidermal wound healing.

Supplementary Video 2 | Live imaging of ER vesicle trafficking.

Supplementary Video 3 | Live imaging of Fat body.

Supplementary Video 4 | Live imaging of trachea.

Supplementary Video 5 | Live imaging of wing imaginal disc.

Supplementary Video 6 | Live imaging of peripheral neurons.

Supplementary Video 7 | Live imaging of a chordotonal organ.

Supplementary Video 8 | Live imaging of midgut enterocytes.

Supplementary Video 9 | Live imaging of midgut interstitial cells.

Supplementary Video 10 | Live imaging of haemocytes.

Supplementary Video 11 | Live imaging of muscle.

Supplementary Video 12 | Single- and multi-cell laser ablation and wound healing process.

Supplementary Video 13 | Dynamic insulin/PIP3/FOXO signalling during wound healing.

Supplementary Video 14 | Effect of reduced insulin receptor signalling.

## Acknowledgements

We are grateful to M. Kakanj for her photographic and graphical support. We thank S. Roth, A. Schauss, F. Papagiannouli, V. Böhm and C. Lesch for critical reading of manuscript, comments and helpful discussions. We thank A. Schauss, P. Zentis and C. Jüngst from the CECAD imaging facility in Cologne (University of Cologne, Cluster of Excellence in Ageing Research) for support and the Bloomington and VDRC Stock Centers for fly strains. This work was supported by grants from the European Regional Development Fund and the German state North Rhine-Westphalia (NRW im Ziel 2) to S.A.E. and L.P., a CMMC grant to M.L. and S.A.E. and an EMBL fellowship to P.K.

## Author contributions

P.K. conceived, designed and developed long-term live imaging and laser ablation protocol for *Drosophila* larvae, performed all experiments and prepared the figures and tables. P.K., M.L. and L.P. analysed and discussed the data and drafted the manuscript. S.A.E. provided input on wound healing analysis and discussed the data.

## Competing interests

The authors declare no competing financial interests.

## Supplementary information

Supplementary information is available for this paper as a PDF file and Video.

## Related link

Key references using this protocol:

Kakanj, P. *et al.* Insulin and TOR signal in parallel through FOXO and S6K to promote epithelial wound healing. *Nat Commun* **7**, 12972 (2016).

Beati, H. *et al.* The adherens junction-associated LIM domain protein Smallish regulates epithelial morphogenesis. *J Cell Biol* **217**, 1079-1095 (2018).

## Supplementary Video legends

**Supplementary Video. 1 | Live imaging of mitochondria during epidermal wound healing.** Live imaging of mitochondria (green) dynamics during epidermal wound healing. Single-cell wound in the dorsal epidermis of *Drosophila* L3 larva expressing mito-GFP (green) and DsRed2-Nuc (magenta) to mark mitochondria and nuclei in the epidermis: *A58>mito-GFP; DsRed2-Nuc (w^1118^; UAS-mito-HA-GFP/+; A58-Gal4, DsRed2-Nuc/+*). Each frame is a merge of 57 planes spaced 0.28 µm apart. Scale bar: 20 µm. Corresponds to **Fig. 7a-b**.

**Supplementary Video. 2 | Live imaging of ER vesicle trafficking.** Live imaging of ER vesicle (green) trafficking between ER and Golgi in the epidermis of *Drosophila* L3 larva expressing *e22c>; RFP-KDEL (w^1118^; e22c-Gal4/+; UASp-RFP.KDEL/+*). KDEL is an ER marker, encodes a sequence to prevent secretion of the protein form ER or retrieval of ER proteins from the Golgi. Each frame is a single plane. Scale bar: 20 µm. Corresponds to **Fig. 7c-d**.

**Supplementary Video. 3 | Live imaging of Fat body.** Live imaging of fat body (green) in the L3 larvae expressing *c7>mCD8-GFP* (*w^1118^; c7-Gal4/+; UAS-mCD8-GFP/+*). The mCD8-GFP marks the cell membranes. Each frame is a single plane. Scale bars: first movie 80 µm and second movie 20 µm. Corresponds to **Fig. 7e-f**.

**Supplementary Video. 4 | Live imaging of trachea. (a-b)** Live imaging of trachea (green) in the early L3 larvae in two individuals both expressing *btl>GFP* (*w^1118^; btl-Gal4, UAS-GFP; +*). Each frame is a merge of 21 planes spaced 0.28 µm apart. Scale bar: (**a,b**) 17 µm. Corresponds to **Fig. 7g-h**.

**Supplementary Video. 5 | Live imaging of wing imaginal disc.** First movie shows a live imaging through the z-stacks of dorsal compartment of wing imaginal disk (grey) in late L3 larva expressing *hh>mCD8-GFP* (*w^1118^; +; hh-Gal4, UAS-mCD8-GFP/+*). The mCD8-GFP marks the cell membranes. Each frame is a single plane of z-stacks (54 stacks with 1µm space). Scale bar: 50 µm. Second movie shows a short live imaging of entire imaginal disk (green) in early L3 larvae expressing *esg>GFP* (*w^1118^; esg-Gal4, UAS-GFP/+; +*). Each frame is a single plane. Scale bar: 100 µm. Corresponds to **Fig. 7i-k**.

**Supplementary Video. 6 | Live imaging of peripheral neurons.** Live imaging of peripheral neuron (black) in the L3 larva expressing mCD8-GFP (black) to mark cell membranes of the neurons: *elav>mCD8-GFP* (*w^1118^, elave-Gal4/UAS-mCD8-GFP; UAS-mCD8-GFP/+; UAS-mCD8-GFP/+*). Each frame is a merge of 21 planes spaced 0.28 µm apart. Scale bar: 15 µm. Corresponds to **Fig. 7m**.

**Supplementary Video. 7 | Live imaging of a chordotonal organ.** Live imaging of lateral pentascolopidial (Lch5) organ (green) in L3 larva expressing mCD8-GFP (green) to mark cell membranes: *elav>mCD8-GFP* (*w^1118^, elave-Gal4/UAS-mCD8-GFP; UAS-mCD8-GFP/+; UAS-mCD8-GFP/+*). The Lch5 organ is a particular chordotonal organ, which plays a role by proprioceptive locomotion control. Each frame is a single plane. Scale bar: 8 µm.

**Supplementary Video. 8 | Live imaging of midgut enterocytes.** Live imaging of midgut enterocytes cells (green) in the L3 larva expressing *NP1>GFP-Atg8a; foxo* (*w^1118^; NP1-Gal4/UAS-GFP-Atg8a; UAS-foxo/+*). Overexpression of FOXO in the midgut enterocytes induces autophagy (green dots are autophagosomes). The fine-focus was changed during the live imaging to image through different planes. Each frame is a single plane. Scale bar: 20 µm. Corresponds to **Fig. 7n**.

**Supplementary Video. 9 | Live imaging of midgut interstitial cells.** Live imaging of midgut interstitial cells (green) in the L3 larva expressing *NP1>GFP-Atg8a; foxo* (*w^1118^; NP1-Gal4/UAS-GFP-Atg8a; UAS-foxo/+*). Overexpression of FOXO induces autophagy in the midgut interstitial cells (green dots are autophagosomes). The fine-focus was changed during the live imaging to image through different planes. Each frame is a single plane. Scale bar: 20 µm. Corresponds to **Fig. 7o**.

**Supplementary Video. 10 | Live imaging of haemocytes.** Live imaging of haemocytes in the L3 larva expressing *Hml>dsRed* (magenta, in the first movie) to mark the entire cytoplasm of the haemocytes and Jupiter (green, in the second movie) to mark the microtubules of the haemocytes: *Hml>dsRed* (*w^1118^; Hml-Gal4/+; UAS-dsRed/+*) and *Jupiter-GFP* trap (*w^1118^; +; Jupiter-GFP*). Each frame is a merge of 21 planes spaced 0.28 µm apart. Scale bars: 20 µm. Corresponds to **Fig. 7p-q**.

**Supplementary Video. 11 | Live imaging of muscle.** First movie shows live imaging of dorsal muscles (green) in L3 larva expressing *MHC>GFP-Atg8a; foxo* (*w^1118^; MHC-Gal4/UAS-GFP-Atg8a; UAS-foxo/+*), in which high level of FOXO induces autophagy (green dots are autophagosomes). Second movie shoes live imaging of dorsal muscles (green) in L3 larva expressing *Nrg-GFP trap (w^1118^, Ngr-GFP; +; +*). In both movies the fine-focus was changed during the live imaging to image through different planes. Each frame is a single plane. Scale bar: 20 µm. Corresponds to **Fig. 7r-s**.

**Supplementary Video. 12 | Single- and multi-cell laser ablation and wound healing process.** Live imaging of single-cell (first movie) and multi-cell (second movie) laser wounding in the dorsal epidermis of L3 larva expressing Src-GFP (green) and DsRed2-Nuc (magenta) to mark cell membrane and nuclei in the epidermis: *A58>Src-GFP, DsRed2-Nuc (w^1118^; +; A58-Gal4, UAS-Src-GFP, UAS-DsRed2-Nuc/+*). Each frame is a merge of 57 planes spaced 0.28 µm apart. Scale bars: 20 µm. Corresponds to **Fig. 10**.

**Supplementary Video. 13 | Dynamic insulin/PIP3/FOXO signalling during wound healing.** Redistribution of PIP3 reporter, tGPH (green) and FOXO (magenta) shuttling after wounding in the cells directly surrounding the wound. Wound healing was performed in the dorsal epidermis of L3 larva expressing *tGPH* (tubulin:GFP-PH) *and foxo-mCherry* (*w^-^; tGPH; endo-dfoxo-v3-mCherry*). Each frame is a merge of 57 planes spaced 0.28 µm apart. Scale bar: 20 µm. Corresponds to **Fig. 11a**.

**Supplementary Video. 14 | Effect of reduced insulin receptor signalling.** Lowering of insulin receptor signalling in the larvae by expressing a dominant negative version of the insulin receptor (InR^DN^), reduced accumulation of the PIP3-reporter tGPH (back) at the wound edges and slows down the wound healing processes. Transgene genotypes: control (*w^1118^; tGPH/+; A58-Gal4/+*) and *A58>InR^DN^* (*w^1118^; tGPH/+; A58-Gal4/UAS-InR^DN^*). Each frame is a merge of 57 planes spaced 0.28 µm apart. Scale bar: 20 µm. Corresponds to **Fig. 11b**.

## References

1. Dobie, K.W. et al. Identification of chromosome inheritance modifiers in Drosophila melanogaster. Genetics 157, 1623–1637 (2001).

2. Hales, K.G., Korey, C.A., Larracuente, A.M. & Roberts, D.M. Genetics on the Fly: A Primer on the Drosophila Model System. Genetics 201, 815–842 (2015).

3. Wieschaus, E. & Nusslein-Volhard, C. The Heidelberg Screen for Pattern Mutants of Drosophila: A Personal Account. Annu Rev Cell Dev Biol 32, 1–46 (2016).

4. Abreu-Blanco, M.T., Verboon, J.M. & Parkhurst, S.M. Cell wound repair in Drosophila occurs through three distinct phases of membrane and cytoskeletal remodeling. J Cell Biol 193, 455–464 (2011).

5. Tsai, C.R., Wang, Y. & Galko, M.J. Crawling wounded: molecular genetic insights into wound healing from Drosophila larvae. Int J Dev Biol 62, 479–489 (2018).

6. Razzell, W., Wood, W. & Martin, P. Swatting flies: modelling wound healing and inflammation in Drosophila. Dis Model Mech 4, 569–574 (2011).

7. Losick, V.P., Fox, D.T. & Spradling, A.C. Polyploidization and cell fusion contribute to wound healing in the adult Drosophila epithelium. Curr Biol 23, 2224–2232 (2013).

8. Belacortu, Y. & Paricio, N. Drosophila as a model of wound healing and tissue regeneration in vertebrates. Dev Dyn 240, 2379–2404 (2011).

9. Dar, A.C., Das, T.K., Shokat, K.M. & Cagan, R.L. Chemical genetic discovery of targets and anti-targets for cancer polypharmacology. Nature 486, 80–84 (2012).

10. Enomoto, M., Siow, C. & Igaki, T. Drosophila As a Cancer Model. Adv Exp Med Biol 1076, 173–194 (2018).

11. Das, T.K. & Cagan, R.L. A Drosophila approach to thyroid cancer therapeutics. Drug Discov Today Technol 10, e65–71 (2013).

12. Sonoshita, M. et al. A whole-animal platform to advance a clinical kinase inhibitor into new disease space. Nat Chem Biol 14, 291–298 (2018).

13. McGurk, L., Berson, A. & Bonini, N.M. Drosophila as an In Vivo Model for Human Neurodegenerative Disease. Genetics 201, 377–402 (2015).

14. Moloney, A., Sattelle, D.B., Lomas, D.A. & Crowther, D.C. Alzheimer’s disease: insights from Drosophila melanogaster models. Trends Biochem Sci 35, 228–235 (2010).

15. Piper, M.D.W. & Partridge, L. Drosophila as a model for ageing. Biochim Biophys Acta Mol Basis Dis 1864, 2707–2717 (2018).

16. Ugur, B., Chen, K. & Bellen, H.J. Drosophila tools and assays for the study of human diseases. Dis Model Mech 9, 235–244 (2016).

17. Kam, Z., Minden, J.S., Agard, D.A., Sedat, J.W. & Leptin, M. Drosophila gastrulation: analysis of cell shape changes in living embryos by three-dimensional fluorescence microscopy. Development 112, 365–370 (1991).

18. Wang, S. & Hazelrigg, T. Implications for bcd mRNA localization from spatial distribution of exu protein in Drosophila oogenesis. Nature 369, 400–403 (1994).

19. Ninov, N. & Martin-Blanco, E. Live imaging of epidermal morphogenesis during the development of the adult abdominal epidermis of Drosophila. Nat Protoc 2, 3074–3080 (2007).

20. Weavers, H., Franz, A., Wood, W. & Martin, P. Long-term In Vivo Tracking of Inflammatory Cell Dynamics Within Drosophila Pupae. J Vis Exp (2018).

21. Fuger, P., Behrends, L.B., Mertel, S., Sigrist, S.J. & Rasse, T.M. Live imaging of synapse development and measuring protein dynamics using two-color fluorescence recovery after photo-bleaching at Drosophila synapses. Nat Protoc 2, 3285–3298 (2007).

22. Martin, J.L. et al. Long-term live imaging of the Drosophila adult midgut reveals real-time dynamics of division, differentiation and loss. Elife 7 (2018).

23. Antunes, M., Pereira, T., Cordeiro, J.V., Almeida, L. & Jacinto, A. Coordinated waves of actomyosin flow and apical cell constriction immediately after wounding. J Cell Biol 202, 365–379 (2013).

24. Weavers, H., Evans, I.R., Martin, P. & Wood, W. Corpse Engulfment Generates a Molecular Memory that Primes the Macrophage Inflammatory Response. Cell 165, 1658–1671 (2016).

25. Larson, D.E., Liberman, Z. & Cagan, R.L. Cellular behavior in the developing Drosophila pupal retina. Mech Dev 125, 223–232 (2008).

26. Weitkunat, M., Brasse, M., Bausch, A.R. & Schnorrer, F. Mechanical tension and spontaneous muscle twitching precede the formation of cross-striated muscle in vivo. Development 144, 1261–1272 (2017).

27. Bosveld, F. et al. Epithelial tricellular junctions act as interphase cell shape sensors to orient mitosis. Nature 530, 495–498 (2016).

28. Sauerwald, J., Backer, W., Matzat, T., Schnorrer, F. & Luschnig, S. Matrix metalloproteinase 1 modulates invasive behavior of tracheal branches during ingression into Drosophila flight muscles. bioRxiv bioRxiv doi: 10.1101/669028 (2019).

29. Nienhaus, U., Aegerter-Wilmsen, T. & Aegerter, C.M. In-vivo imaging of the Drosophila wing imaginal disc over time: novel insights on growth and boundary formation. PLoS One 7, e47594 (2012).

30. Ghannad-Rezaie, M., Wang, X., Mishra, B., Collins, C. & Chronis, N. Microfluidic chips for in vivo imaging of cellular responses to neural injury in Drosophila larvae. PLoS One 7, e29869 (2012).

31. Heemskerk, I., Lecuit, T. & LeGoff, L. Dynamic clonal analysis based on chronic in vivo imaging allows multiscale quantification of growth in the Drosophila wing disc. Development 141, 2339–2348 (2014).

32. Andlauer, T.F. & Sigrist, S.J. In vivo imaging of the Drosophila larval neuromuscular junction. Cold Spring Harb Protoc 2012, 481–489 (2012).

33. Cevik, D. et al. Chloroform and desflurane immobilization with recovery of viable Drosophila larvae for confocal imaging. J Insect Physiol 117, 103900 (2019).

34. Kakanj, P. et al. Insulin and TOR signal in parallel through FOXO and S6K to promote epithelial wound healing. Nat Commun 7, 12972 (2016).

35. Beati, H. et al. The adherens junction-associated LIM domain protein Smallish regulates epithelial morphogenesis. J Cell Biol 217, 1079–1095 (2018).

36. Griffin, R., Binari, R. & Perrimon, N. Genetic odyssey to generate marked clones in Drosophila mosaics. Proc Natl Acad Sci U S A 111, 4756–4763 (2014).

37. Germani, F., Bergantinos, C. & Johnston, L.A. Mosaic Analysis in Drosophila. Genetics 208, 473–490 (2018).

38. Makhijani, K. et al. Precision Optogenetic Tool for Selective Single- and Multiple-Cell Ablation in a Live Animal Model System. Cell Chem Biol 24, 110–119 (2017).

39. Stein, D.S. & Stevens, L.M. Maternal control of the Drosophila dorsal-ventral body axis. Wiley Interdiscip Rev Dev Biol 3, 301–330 (2014).

40. Tepass, U. et al. shotgun encodes Drosophila E-cadherin and is preferentially required during cell rearrangement in the neurectoderm and other morphogenetically active epithelia. Genes Dev 10, 672–685 (1996).

41. Cox, R.T., Kirkpatrick, C. & Peifer, M. Armadillo is required for adherens junction assembly, cell polarity, and morphogenesis during Drosophila embryogenesis. J Cell Biol 134, 133–148 (1996).

42. Uemura, T. et al. Zygotic Drosophila E-cadherin expression is required for processes of dynamic epithelial cell rearrangement in the Drosophila embryo. Genes Dev 10, 659–671 (1996).

43. Kuhn, H., Sopko, R., Coughlin, M., Perrimon, N. & Mitchison, T. The Atg1-Tor pathway regulates yolk catabolism in Drosophila embryos. Development 142, 3869–3878 (2015).

44. Mirth, C.K. & Shingleton, A.W. Integrating body and organ size in Drosophila: recent advances and outstanding problems. Front Endocrinol (Lausanne) 3, 49 (2012).

45. Vollmer, J., Casares, F. & Iber, D. Growth and size control during development. Open Biol 7 (2017).

46. Johnston, L.A. & Gallant, P. Control of growth and organ size in Drosophila. Bioessays 24, 54–64 (2002).

47. Nijhout, H.F. The control of growth. Development 130, 5863–5867 (2003).

48. Badouel, C. & McNeill, H. Apical junctions and growth control in Drosophila. Biochim Biophys Acta 1788, 755–760 (2009).

49. Diaz-de-la-Loza, M.D. et al. Apical and Basal Matrix Remodeling Control Epithelial Morphogenesis. Dev Cell 46, 23–39 e25 (2018).

50. Tumaneng, K., Russell, R.C. & Guan, K.L. Organ size control by Hippo and TOR pathways. Curr Biol 22, R368–379 (2012).

51. Beira, J.V. & Paro, R. The legacy of Drosophila imaginal discs. Chromosoma 125, 573–592 (2016).

52. Baena-Lopez, L.A., Nojima, H. & Vincent, J.P. Integration of morphogen signalling within the growth regulatory network. Curr Opin Cell Biol 24, 166–172 (2012).

53. Miles, W.O., Dyson, N.J. & Walker, J.A. Modeling tumor invasion and metastasis in Drosophila. Dis Model Mech 4, 753–761 (2011).

54. Stefanatos, R.K. & Vidal, M. Tumor invasion and metastasis in Drosophila: a bold past, a bright future. J Genet Genomics 38, 431–438 (2011).

55. Das, T.K. & Cagan, R.L. Non-mammalian models of multiple endocrine neoplasia type 2. Endocr Relat Cancer 25, T91–T104 (2018).

56. Rudrapatna, V.A., Cagan, R.L. & Das, T.K. Drosophila cancer models. Dev Dyn 241, 107–118 (2012).

57. Bangi, E., Garza, D. & Hild, M. In vivo analysis of compound activity and mechanism of action using epistasis in Drosophila. J Chem Biol 4, 55–68 (2011).

58. Willoughby, L.F. et al. An in vivo large-scale chemical screening platform using Drosophila for anti-cancer drug discovery. Dis Model Mech 6, 521–529 (2013).

59. Gasque, G., Conway, S., Huang, J., Rao, Y. & Vosshall, L.B. Small molecule drug screening in Drosophila identifies the 5HT2A receptor as a feeding modulation target. Sci Rep 3, srep02120 (2013).

60. Burra, S., Wang, Y., Brock, A.R. & Galko, M.J. Using Drosophila larvae to study epidermal wound closure and inflammation. Methods Mol Biol 1037, 449–461 (2013).

61. Galko, M.J. & Krasnow, M.A. Cellular and genetic analysis of wound healing in Drosophila larvae. PLoS Biol 2, E239 (2004).

62. Mishra, B. et al. Using microfluidics chips for live imaging and study of injury responses in Drosophila larvae. J Vis Exp, e50998 (2014).

63. Zhang, Y. et al. In vivo imaging of intact Drosophila larvae at sub-cellular resolution. J Vis Exp (2010).

64. Rasse, T.M. et al. Glutamate receptor dynamics organizing synapse formation in vivo. Nat Neurosci 8, 898–905 (2005).

65. Zirin, J. et al. Ecdysone signaling at metamorphosis triggers apoptosis of Drosophila abdominal muscles. Dev Biol 383, 275–284 (2013).

66. Klionsky, D.J. et al. Guidelines for the use and interpretation of assays for monitoring autophagy (3rd edition). Autophagy 12, 1–222 (2016).

67. Takats, S., Varga, A., Pircs, K. & Juhasz, G. Loss of Drosophila Vps16A enhances autophagosome formation through reduced Tor activity. Autophagy 11, 1209–1215 (2015).

68. Brand, A.H. & Perrimon, N. Targeted gene expression as a means of altering cell fates and generating dominant phenotypes. Development 118, 401–415 (1993).

69. Wang, Y. et al. Integrin Adhesions Suppress Syncytium Formation in the Drosophila Larval Epidermis. Curr Biol 25, 2215–2227 (2015).

70. Lawrence, P.A., Bodmer, R. & Vincent, J.P. Segmental patterning of heart precursors in Drosophila. Development 121, 4303–4308 (1995).

71. Huang, J., Zhou, W., Dong, W., Watson, A.M. & Hong, Y. From the Cover: Directed, efficient, and versatile modifications of the Drosophila genome by genomic engineering. Proc Natl Acad Sci U S A 106, 8284–8289 (2009).

72. Wodarz, A., Hinz, U., Engelbert, M. & Knust, E. Expression of crumbs confers apical character on plasma membrane domains of ectodermal epithelia of Drosophila. Cell 82, 67–76 (1995).

73. Cohen, E. Chitin synthesis and inhibition: a revisit. Pest Manag Sci 57, 946–950 (2001).

74. Maimon, I. & Gilboa, L. Dissection and staining of Drosophila larval ovaries. J Vis Exp (2011).

75. Forster, D. & Luschnig, S. Src42A-dependent polarized cell shape changes mediate epithelial tube elongation in Drosophila. Nat Cell Biol 14, 526–534 (2012).

76. Nehme, N.T. et al. A model of bacterial intestinal infections in Drosophila melanogaster. PLoS Pathog 3, e173 (2007).

77. Jiang, H. et al. Cytokine/Jak/Stat signaling mediates regeneration and homeostasis in the Drosophila midgut. Cell 137, 1343–1355 (2009).

78. Zeng, X., Chauhan, C. & Hou, S.X. Characterization of midgut stem cell- and enteroblast-specific Gal4 lines in drosophila. Genesis 48, 607–611 (2010).

79. Micchelli, C.A. & Perrimon, N. Evidence that stem cells reside in the adult Drosophila midgut epithelium. Nature 439, 475–479 (2006).

80. Lin, D.M. & Goodman, C.S. Ectopic and increased expression of Fasciclin II alters motoneuron growth cone guidance. Neuron 13, 507–523 (1994).

81. Torroja, L., Packard, M., Gorczyca, M., White, K. & Budnik, V. The Drosophila beta-amyloid precursor protein homolog promotes synapse differentiation at the neuromuscular junction. J Neurosci 19, 7793–7803 (1999).

82. Gronke, S. et al. Control of fat storage by a Drosophila PAT domain protein. Curr Biol 13, 603–606 (2003).

83. Goto, A., Kadowaki, T. & Kitagawa, Y. Drosophila hemolectin gene is expressed in embryonic and larval hemocytes and its knock down causes bleeding defects. Dev Biol 264, 582–591 (2003).

84. Stramer, B. et al. Live imaging of wound inflammation in Drosophila embryos reveals key roles for small GTPases during in vivo cell migration. J Cell Biol 168, 567–573 (2005).

85. Lee, S. & Kolodziej, P.A. The plakin Short Stop and the RhoA GTPase are required for E-cadherin-dependent apical surface remodeling during tracheal tube fusion. Development 129, 1509–1520 (2002).

86. Ranganayakulu, G., Schulz, R.A. & Olson, E.N. Wingless signaling induces nautilus expression in the ventral mesoderm of the Drosophila embryo. Dev Biol 176, 143–148 (1996).

87. Klein, P. et al. Ret rescues mitochondrial morphology and muscle degeneration of Drosophila Pink1 mutants. EMBO J 33, 341–355 (2014).

88. Kaltschmidt, J.A., Davidson, C.M., Brown, N.H. & Brand, A.H. Rotation and asymmetry of the mitotic spindle direct asymmetric cell division in the developing central nervous system. Nat Cell Biol 2, 7–12 (2000).

89. Lee, T. & Luo, L. Mosaic analysis with a repressible cell marker for studies of gene function in neuronal morphogenesis. Neuron 22, 451–461 (1999).

90. Pfeiffer, B.D. et al. Refinement of tools for targeted gene expression in Drosophila. Genetics 186, 735–755 (2010).

91. Morin, X., Daneman, R., Zavortink, M. & Chia, W. A protein trap strategy to detect GFP-tagged proteins expressed from their endogenous loci in Drosophila. Proc Natl Acad Sci U S A 98, 15050–15055 (2001).

92. Oda, H. & Tsukita, S. Real-time imaging of cell-cell adherens junctions reveals that Drosophila mesoderm invagination begins with two phases of apical constriction of cells. J Cell Sci 114, 493–501 (2001).

93. Cai, D. et al. Mechanical feedback through E-cadherin promotes direction sensing during collective cell migration. Cell 157, 1146–1159 (2014).

94. Royou, A., Sullivan, W. & Karess, R. Cortical recruitment of nonmuscle myosin II in early syncytial Drosophila embryos: its role in nuclear axial expansion and its regulation by Cdc2 activity. J Cell Biol 158, 127–137 (2002).

95. Martin, A.C., Kaschube, M. & Wieschaus, E.F. Pulsed contractions of an actin-myosin network drive apical constriction. Nature 457, 495–499 (2009).

96. Buszczak, M. et al. The carnegie protein trap library: a versatile tool for Drosophila developmental studies. Genetics 175, 1505–1531 (2007).

97. Harris, T.J. & Peifer, M. Adherens junction-dependent and -independent steps in the establishment of epithelial cell polarity in Drosophila. J Cell Biol 167, 135–147 (2004).

98. Barolo, S., Castro, B. & Posakony, J.W. New Drosophila transgenic reporters: insulated P-element vectors expressing fast-maturing RFP. Biotechniques 36, 436–440, 442 (2004).

99. Polychronidou, M., Hellwig, A. & Grosshans, J. Farnesylated nuclear proteins Kugelkern and lamin Dm0 affect nuclear morphology by directly interacting with the nuclear membrane. Mol Biol Cell 21, 3409–3420 (2010).

100. Katsani, K.R., Karess, R.E., Dostatni, N. & Doye, V. In vivo dynamics of Drosophila nuclear envelope components. Mol Biol Cell 19, 3652–3666 (2008).

101. Feng, K., Palfreyman, M.T., Hasemeyer, M., Talsma, A. & Dickson, B.J. Ascending SAG neurons control sexual receptivity of Drosophila females. Neuron 83, 135–148 (2014).

102. Okajima, T., Xu, A., Lei, L. & Irvine, K.D. Chaperone activity of protein O-fucosyltransferase 1 promotes notch receptor folding. Science 307, 1599–1603 (2005).

103. Okajima, T., Reddy, B., Matsuda, T. & Irvine, K.D. Contributions of chaperone and glycosyltransferase activities of O-fucosyltransferase 1 to Notch signaling. BMC Biol 6, 1 (2008).

104. Pilling, A.D., Horiuchi, D., Lively, C.M. & Saxton, W.M. Kinesin-1 and Dynein are the primary motors for fast transport of mitochondria in Drosophila motor axons. Mol Biol Cell 17, 2057–2068 (2006).

105. Bardet, P.L. et al. A fluorescent reporter of caspase activity for live imaging. Proc Natl Acad Sci U S A 105, 13901–13905 (2008).

106. Nezis, I.P. et al. Autophagic degradation of dBruce controls DNA fragmentation in nurse cells during late Drosophila melanogaster oogenesis. J Cell Biol 190, 523–531 (2010).

107. Arsham, A.M. & Neufeld, T.P. A genetic screen in Drosophila reveals novel cytoprotective functions of the autophagy-lysosome pathway. PLoS One 4, e6068 (2009).

108. Rusten, T.E. et al. Programmed autophagy in the Drosophila fat body is induced by ecdysone through regulation of the PI3K pathway. Dev Cell 7, 179–192 (2004).

109. Chang, Y.Y. & Neufeld, T.P. An Atg1/Atg13 complex with multiple roles in TOR-mediated autophagy regulation. Mol Biol Cell 20, 2004–2014 (2009).

110. Hatan, M., Shinder, V., Israeli, D., Schnorrer, F. & Volk, T. The Drosophila blood brain barrier is maintained by GPCR-dependent dynamic actin structures. J Cell Biol 192, 307–319 (2011).

111. Grieder, N.C., de Cuevas, M. & Spradling, A.C. The fusome organizes the microtubule network during oocyte differentiation in Drosophila. Development 127, 4253–4264 (2000).

112. Megraw, T.L., Kilaru, S., Turner, F.R. & Kaufman, T.C. The centrosome is a dynamic structure that ejects PCM flares. J Cell Sci 115, 4707–4718 (2002).

113. Zirin, J., Nieuwenhuis, J. & Perrimon, N. Role of autophagy in glycogen breakdown and its relevance to chloroquine myopathy. PLoS Biol 11, e1001708 (2013).

114. Klapholz, B. et al. Alternative mechanisms for talin to mediate integrin function. Curr Biol 25, 847–857 (2015).

115. Benton, R. & St Johnston, D. A conserved oligomerization domain in drosophila Bazooka/PAR-3 is important for apical localization and epithelial polarity. Curr Biol 13, 1330–1334 (2003).

116. Khuong, T.M., Habets, R.L., Slabbaert, J.R. & Verstreken, P. WASP is activated by phosphatidylinositol-4,5-bisphosphate to restrict synapse growth in a pathway parallel to bone morphogenetic protein signaling. Proc Natl Acad Sci U S A 107, 17379–17384 (2010).

117. Britton, J.S., Lockwood, W.K., Li, L., Cohen, S.M. & Edgar, B.A. Drosophila’s insulin/PI3-kinase pathway coordinates cellular metabolism with nutritional conditions. Dev Cell 2, 239–249 (2002).

118. Karpova, N., Bobinnec, Y., Fouix, S., Huitorel, P. & Debec, A. Jupiter, a new Drosophila protein associated with microtubules. Cell Motil Cytoskeleton 63, 301–312 (2006).

